# A Brainwide Atlas of Synaptic Nanoarchitecture Across the Mouse Lifespan

**DOI:** 10.64898/2026.02.18.706606

**Authors:** Takeshi Kaizuka, Zhen Qiu, Edita Bulovaite, Katie Morris, Tianxiao Zhao, Candace Adams, Gabor Varga, Digin Dominic, Noboru H. Komiyama, Mathew H. Horrocks, Seth G.N. Grant

**Affiliations:** Genes to Cognition Programme, Institute for Neuroscience and Cardiovascular Research, University of Edinburgh, Edinburgh EH16 4SB, UK; Department of Biomedical Engineering, University of Strathclyde, Glasgow G4 0NW, UK; School of Chemistry, University of Edinburgh, David Brewster Road, Edinburgh EH9 3FJ, UK; IRR Chemistry Hub, Institute for Regeneration and Repair, University of Edinburgh, Edinburgh EH16 4UU, UK; Simons Initiative for the Developing Brain (SIDB), Institute for Neuroscience and Cardiovascular Research, University of Edinburgh, Edinburgh EH8 9XD, UK; The Patrick Wild Centre for Research into Autism, Fragile X Syndrome & Intellectual Disabilities, Institute for Neuroscience and Cardiovascular Research, University of Edinburgh, Edinburgh EH8 9XD, UK; Muir Maxwell Epilepsy Centre, Institute for Neuroscience and Cardiovascular Research, University of Edinburgh, Edinburgh EH8 9XD, UK; Euan MacDonald Centre, University of Edinburgh, Edinburgh EH16 4SB, UK

## Abstract

How biological complexity emerges from the ordered assembly of molecular building blocks into supramolecular systems remains a central question, particularly in the mammalian brain with its vast synaptic diversity. We introduce NanoSYNMAP, a genetic, optical, and computational platform that integrates FRET with synaptome mapping to quantify nanoscale proximity of proteins in individual synapses brain-wide. We generate the first brain atlas of synaptic nanoarchitecture, based on the proximity of postsynaptic MAGUK supercomplexes. This reveals a molecular logic in which spacing of supramolecular assemblies specifies nanoscale architecture that organizes the global synaptome architecture. Nanoarchitecture varies across brain regions, differentiates during postnatal development, and remodels with aging. Supercomplex proximity reflects scaffold abundance, nanodomain organization, and competitive interactions among MAGUK assemblies. Deletion of a neuropsychiatric risk gene triggers widespread reorganization of nanoscale architecture. These findings establish molecular proximity as a fundamental scalable dimension of synapse diversity in health and disease.

## INTRODUCTION

Biological complexity emerges from the ordered assembly of molecular building blocks into supramolecular systems. Defining how these assemblies are constructed and remodeled across development, aging, and disease is fundamental to understanding how complex tissues and organs function. Nowhere is this more formidable than in the mammalian brain, where immense complexity arises from the combinatorial molecular diversity of synapses. Each synapse contains thousands of proteins (1–3) that are differentially distributed to generate a vast repertoire of synapse types collectively termed the synaptome (4–8). These synapses are organized into a three-dimensional synaptome architecture that supports the physiological and behavioral operations of brain networks. Across the lifespan, this architecture is dynamically reconfigured (5), depends on sleep for stability (7), and is altered by experience (9), disease-causing mutations (4, 6, 8) and traumatic injury (10), reflecting the interplay between intrinsic genetic programs and environmental influences.

A central determinant of the molecular organization of synapses is the supramolecular organization of synaptic proteins into complexes and supercomplexes (4, 11–13). In the postsynaptic proteome, the most abundant supercomplexes are built around the membrane-associated guanylate kinase (MAGUK) family of scaffolding proteins comprising PSD95 (Dlg4), SAP102 (Dlg3), and PSD93 (Dlg2) (14). They form distinct, biochemically-separable assemblies that tether neurotransmitter receptors, ion channels, adhesion molecules, enzymes, and cytoskeletal elements (3, 11, 15–19). Biochemical single-molecule imaging studies reveal that PSD95 supercomplexes contain two copies of PSD95 (13, 20), and super-resolution microscopy (∼40 nm resolution) shows that PSD95 is organized into subsynaptic nanodomains (also known as “nanoclusters”) and disc-shaped assemblies beneath the postsynaptic membrane (21–24). These nanostructures align with presynaptic release machinery via nanocolumns, highlighting the critical role of supramolecular positioning in determining synaptic efficacy and plasticity (25–30).

Characterizing the nanoscale physical organization of supercomplexes in individual synapses in brain tissue and determining whether this level of molecular organization underpins synaptic diversity and synaptome architecture at the brainwide scale pose substantial technical challenges. Ideally, the proximity between complexes would be measured at resolutions below 10 nm (comparable to the size of an AMPA receptor). Yet, such resolution is currently beyond the reach of most super-resolution methods, and these approaches are not easily scalable to the systematic imaging of millions of synapses across all brain regions. Here, we report the development of a comprehensive toolbox of reagents, methods, and computational approaches for studying the proximity of supercomplexes within synapses across the whole brain and lifespan, in both healthy and diseased mice. These tools are integrated into a nanoscale synaptome mapping (NanoSYNMAP) pipeline, which integrates Förster resonance energy transfer (FRET) – a powerful method for evaluating the proximity of molecules – with our synaptome mapping pipeline. By evaluating the degree of proximity between PSD95 supercomplexes in excitatory synapses across the mouse brain, we uncover nanoscale synapse diversity that is spatially and temporally organized into a global synaptome architecture – the nanoscale synaptome architecture (NSA). We further show that the interdependence between MAGUK supercomplexes governs the brainwide synaptome architecture, providing both a mechanistic basis for the action of psychiatric disease risk genes and a principle for generating synapse diversity. In addition to addressing a major gap in our understanding of brain molecular organization, the methods developed here have wide application for studying the molecular organizing principles of any tissue.

## RESULTS

We developed a suite of tools to label endogenous synaptic proteins and evaluate their physical proximity within individual protein complexes and within synapses (Fig. 1A). These tools include mouse lines in which genetically encoded tags are fused to the C-terminus of endogenous MAGUK proteins (PSD95 and SAP102), enabling labeling with either the self-labeling tag, HaloTag (8) or fluorescent proteins (eGFP or mKO2) (4). In addition, we developed a mouse line in which the C-terminus of PSD95 is labeled with CLIP-tag (Fig. S1). The covalent binding capacity of HaloTag and CLIP-tag for diverse exogenous ligands, together with the spectral properties of eGFP and mKO2, enables the use of multiple fluorophore combinations. These versatile tools allowed us to analyze individual complexes at single-molecule resolution and to evaluate the proximity between complexes within synapses in both synaptosome preparations and tissue sections.

**Figure 1.**
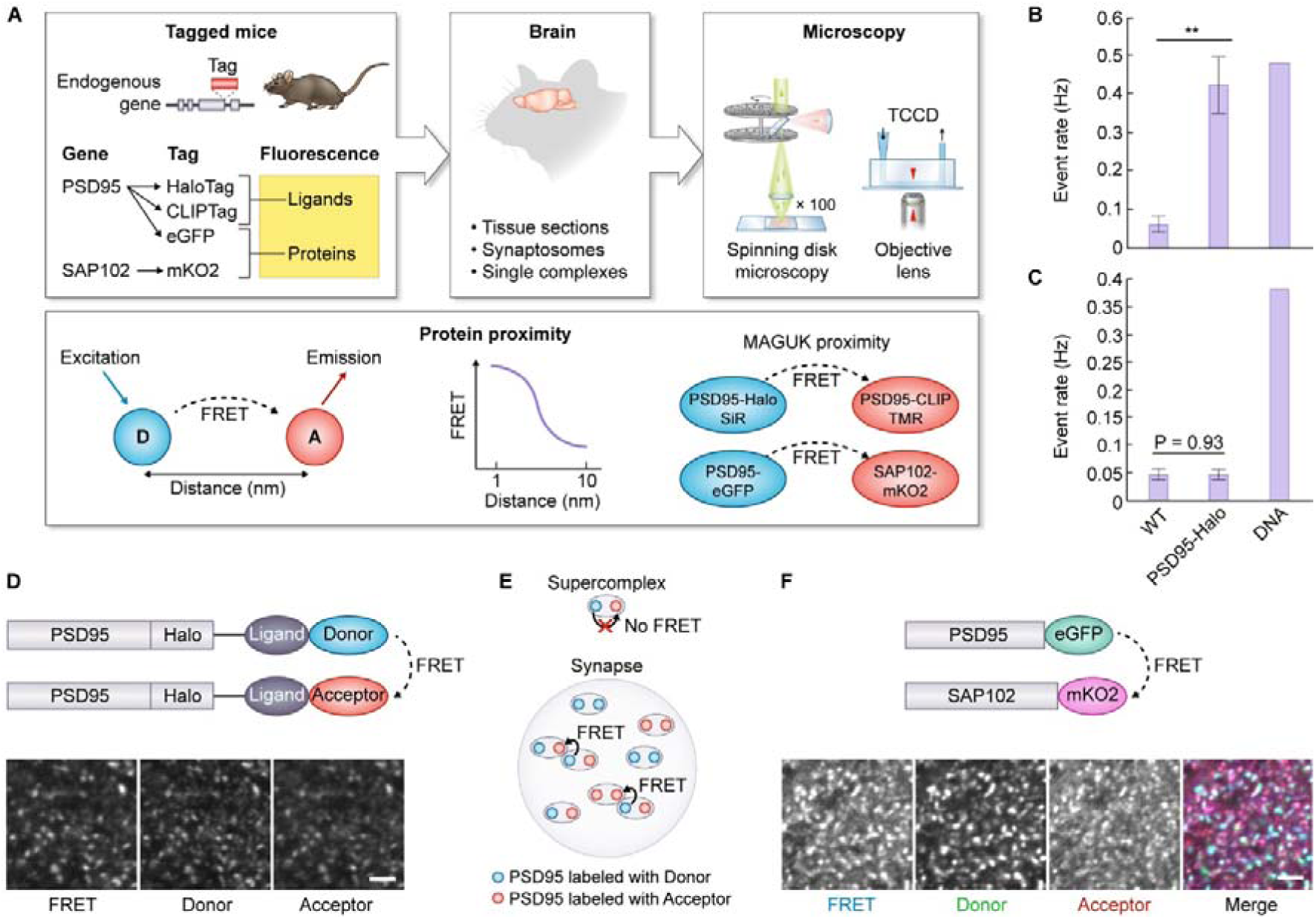
Evaluation of MAGUK proximity in complexes and synapses using NanoSYNMAP. (**A**) The experimental workflow of NanoSYNMAP. Genetically modified mice in which the postsynaptic scaffold proteins PSD95 and/or SAP102 are tagged with a self-labeling tag (HaloTag or CLIP-tag) or a fluorescent protein were used. Brain tissue sections, synaptosomes, or single protein complexes were prepared from the mice. PSD95 carrying a self-labeling tag was labeled with fluorophore-conjugated ligands. The samples were then analyzed by spinning disk confocal microscopy. Individual supercomplexes were studied using single-molecule confocal microscopy: 2-color excitation allowed for the detection of all supercomplexes, whereas 1-color excitation of the donor molecule gave rise to coincident signal in only those supercomplexes where the donor and acceptor dyes were close enough for FRET to occur. (**B**,**C**) Quantification of the coincident event rates showed that whereas a DNA duplex in which the fluorophores were separated by 2 nm (DNA) gave rise to events in both the 2-color (B) and 1-color (C) regimes, the supercomplexes derived from PSD95-HaloTag mice (PSD95-HaloTag) only demonstrated coincident signal in the 2-color case (B). Supercomplexes derived from wild-type mouse (WT) were not detected with either excitation. Data show mean ± SD (n = 3). **P < 0.01, Welch’s t-test. (**D**) Detection of FRET between donor and acceptor fluorophores on synapses in hippocampal CA1sr region. Sections obtained from PSD95–HaloTag mouse were labeled with HaloTag ligands conjugated with donor, acceptor, or a mixture. The fluorescence of the donor, acceptor, and FRET between them was detected using spinning disk confocal microscopy. Scale bar: 2 µm. See also Fig. S2A. (**E**) The postsynaptic proteome model. The observed FRET arises not from PSD95–PSD95 interactions within a single supercomplex, but from the close apposition of PSD95 molecules residing in distinct supercomplexes. (**F**) Detection of FRET between PSD95-eGFP (donor) and SAP102-mKO2 (acceptor) on synapses in hippocampal CA1sr region. Scale bar: 2 µm. See also Fig. S2C.

### FRET does not occur within PSD95 supercomplexes

Biochemical (13) and single-molecule imaging (20) studies have shown that PSD95 supercomplexes typically contain two copies of PSD95, raising the possibility of FRET occurring between fluorophore-tagged versions of the protein. However, our previous work demonstrated that fluorophores genetically tagged to PSD95 are separated by ∼12.7 nm (20), which is beyond the Förster radius of most fluorophore pairs (31). This separation makes FRET within a single supercomplex unlikely. To test this directly, we isolated PSD95–HaloTag complexes from brain homogenates (20), labeled them with FRET donor- and acceptor-conjugated ligands, and examined them using single-molecule confocal microscopy (32) with both one-color and two-color excitation. With two-color excitation, both fluorophores within a supercomplex were excited and detected using two-color coincidence detection (TCCD) (33, 34), regardless of their separation distance. In the one-color excitation setup, only the donor is excited; emission from the acceptor occurs only if FRET is possible (35), implying a donor–acceptor separation of less than 10 nm. Comparing event counts from one-color and two-color excitation allows estimation of the fraction of complexes that are close enough for FRET to occur. We previously used this approach to quantify the proximity of ubiquitin within dimers (36), and here validated the method using a DNA duplex with a 2 nm donor–acceptor separation, which generated signals in both excitation conditions. As expected, PSD95–HaloTag complexes showed significantly more events in TCCD compared with wild-type extracts (Fig. 1B), but FRET-positive events were very rare in either sample (Fig. 1C). This indicates that FRET does not occur within individual PSD95 complexes, consistent with the measured separation distance between fluorophores.

### FRET between MAGUK supercomplexes within synapses

Having shown that FRET does not occur between PSD95 molecules within supercomplexes, we next sought to determine whether FRET was possible between PSD95 in neighboring supercomplexes within the intact synapse. Brain tissue sections from PSD95–HaloTag mice were labeled with HaloTag ligands conjugated to the donor Janelia Fluor 552 (JF552) and the acceptor Janelia Fluor X 650 (JFX650) (Figs. 1D and S2A). Spinning disk confocal microscopy of the hippocampal CA1sr region was performed using donor excitation (561 nm), yielding puncta that were detectable in both the donor and acceptor channels. Single-label controls were included to quantify donor-to-acceptor crosstalk and direct acceptor excitation (see Fig. S2A). Similarly, FRET was detected in mice co-expressing PSD95–HaloTag and PSD95–CLIP-tag labeled with silicon rhodamine (SiR) and CLIP-tag with tetramethylrhodamine (TMR), respectively (Fig. S2B). These findings suggest that within synapses, fluorophores attached to the C-terminus of PSD95 on neighboring supercomplexes can approach within the Förster radius (estimated at 4.2 nm for JF552/JFX650; see Methods) and therefore can be closer than the ∼12.7 nm separation measured for fluorophores within individual PSD95 supercomplexes.

The most straightforward explanation for this observation is that certain PSD95 supercomplexes are closely packed together within synapses (Fig. 1E). To investigate whether a similar proximity also occurs between different types of MAGUK family supercomplexes, we examined the potential for FRET between PSD95 and SAP102. These two MAGUK proteins assemble into distinct supercomplexes of approximately 1.5 MDa and 350 kDa, respectively (11), and are both co-expressed in the Type 3 (PSD95+ SAP102+) subpopulation of excitatory synapses (4). We analyzed brain sections from mice co-expressing PSD95-eGFP and SAP102-mKO2 and observed FRET between PSD95-eGFP and SAP102-mKO2 in hippocampal synapses (Figs. 1F and S2C). These results demonstrate that distinct MAGUK supercomplexes can be in very close proximity within the synapse. For clarity, we refer to the FRET-based evaluation of proximity between PSD95 supercomplexes as the PSD95–PSD95 supercomplex proximity index (P/P-SPI), and that between PSD95 and SAP102 supercomplexes as the PSD95–SAP102 supercomplex proximity index (P/S-SPI). In both cases, higher index values indicate closer physical association between the respective supercomplexes.

### A brainwide synaptome architecture of supercomplex proximity

We next asked whether synapses exhibiting different P/P-SPI values are spatially organized across overarching brain areas and subregions. We developed a pipeline that incorporates FRET measurements into synaptome architecture. This pipeline uses spinning disk confocal microscopy to image individual synapses across whole-brain sections and computational processing to detect and quantify multiple molecular parameters of each synapse. The resulting pipeline is referred to as NanoSYNMAP.

As a first step, we asked whether P/P-SPI and P/S-SPI vary across synapses. In hippocampal CA1sr of PSD95–HaloTag mice, P/P-SPI ranged from 10–40% and showed a broad distribution, which was approximated by a single Gaussian with a full width at half maximum of 16.3% and centered at 26.0% (Fig. 2A). A similar distribution was observed in experiments using PSD95–HaloTag/CLIP-tag mice (Fig. S3). The P/S-SPI ranged from 25–48%, indicating that synaptic variation occurred between supercomplexes built from different scaffold proteins (Fig. 2B). Synapse variation in P/P-SPI and P/S-SPI was also observed in isolated synaptosomes (Figs. 2C,D). However, the values were lower than in tissue and the distributions appeared bimodal and did not adequately fit a single Gaussian distribution, likely reflecting increased intermolecular distances caused by loosening of protein packing during biochemical isolation. These findings indicate that the synaptome is composed of synapses that differ in their nanoscale organization of supercomplexes.

**Figure 2.**
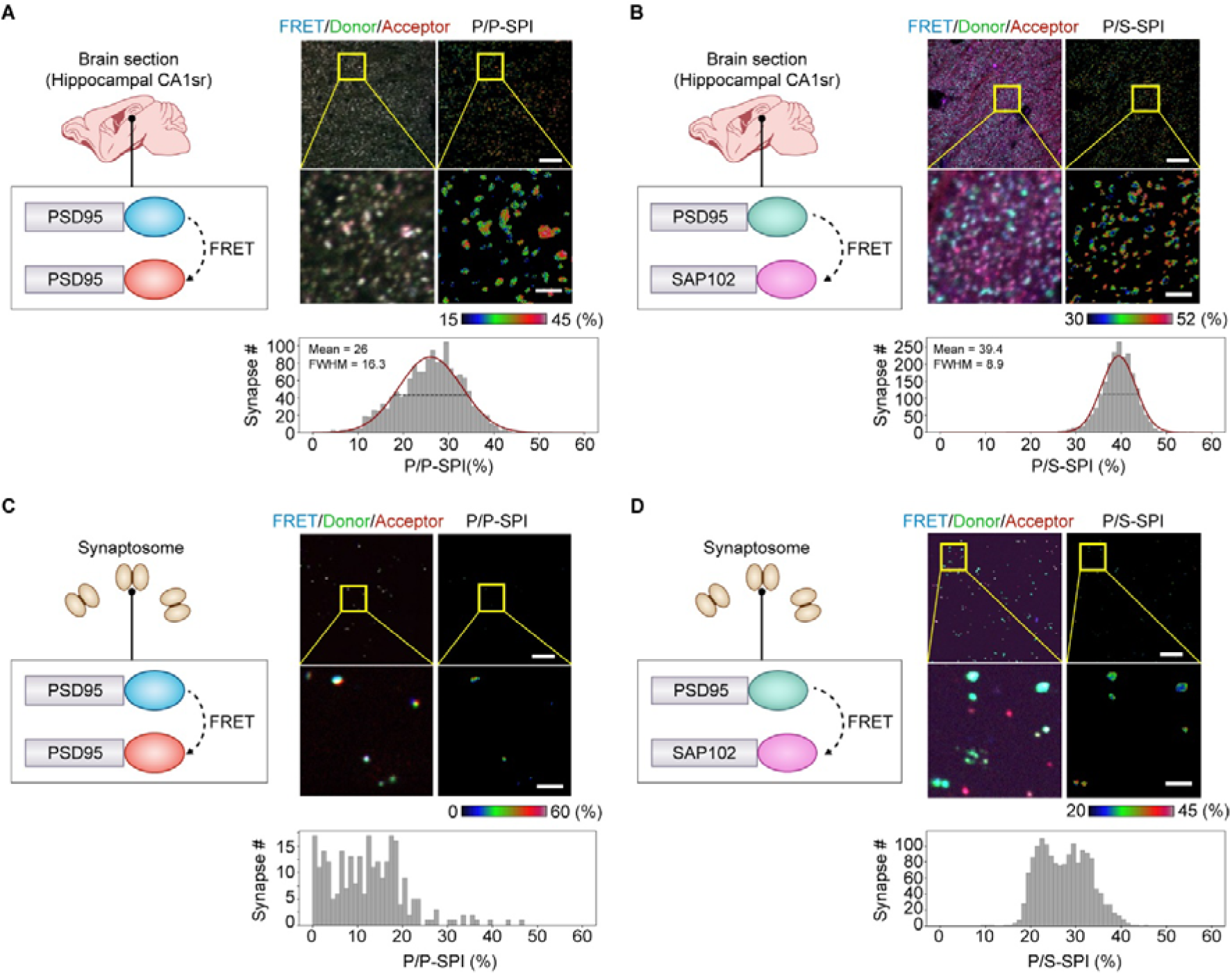
Diversity of synaptic nanoarchitecture in hippocampal CA1sr region. (**A**,**B**) Visualization of synaptic SPI between MAGUK proteins in hippocampal CA1sr region. Sections from PSD95–HaloTag mouse labeled with a mixture of HaloTag ligands conjugated with donor and acceptor fluorophores (A) or sections from PSD95-eGFP SAP102-mKO2 mouse (B) were imaged using spinning disk confocal microscopy. Merged image of the donor, acceptor, and FRET channels (left panels) and heatmap image of SPI (right panels), and histogram of SPI of individual synaptic puncta (bottom panel) are shown. The Gaussian fit to the distribution (red line), together with its mean value and full width at half maximum (FWHM; black dashed line), is shown on the histogram. (**C**,**D**) Visualization of SPI between MAGUK proteins in biochemically-isolated synaptosomes. Synaptosome fraction isolated from forebrain homogenate of PSD95–HaloTag mouse labeled with HaloTag ligands (C) or that of PSD95-eGFP SAP102-mKO2 mouse (D) were imaged and analyzed. Scale bars (A-D): 10 µm (top) and 2 µm (bottom).

Having established that nanoscale packing of MAGUK complexes varies among synapses in hippocampal CA1sr, we used NanoSYNMAP to map entire sagittal sections of 4-month-old adult mice (Fig. 3A) and calculated the P/P-SPI in ∼10 million synapses per section across 112 brain subregions registered to the Allen Brain Atlas (Supplementary Table 1). Representative images of whole-brain sections (Fig. 3A) and hippocampal formation (HPF) (Fig. 3B) clearly show regional differences in P/P-SPI. For each subregion across six mice, we calculated the mean P/P-SPI and generated a brain atlas of nanoscale synaptic organization (Figs. 3C,D). These findings reveal that synapses with distinct nanoscale molecular architectures are spatially organized, producing the NSA of the brain.

**Figure 3.**
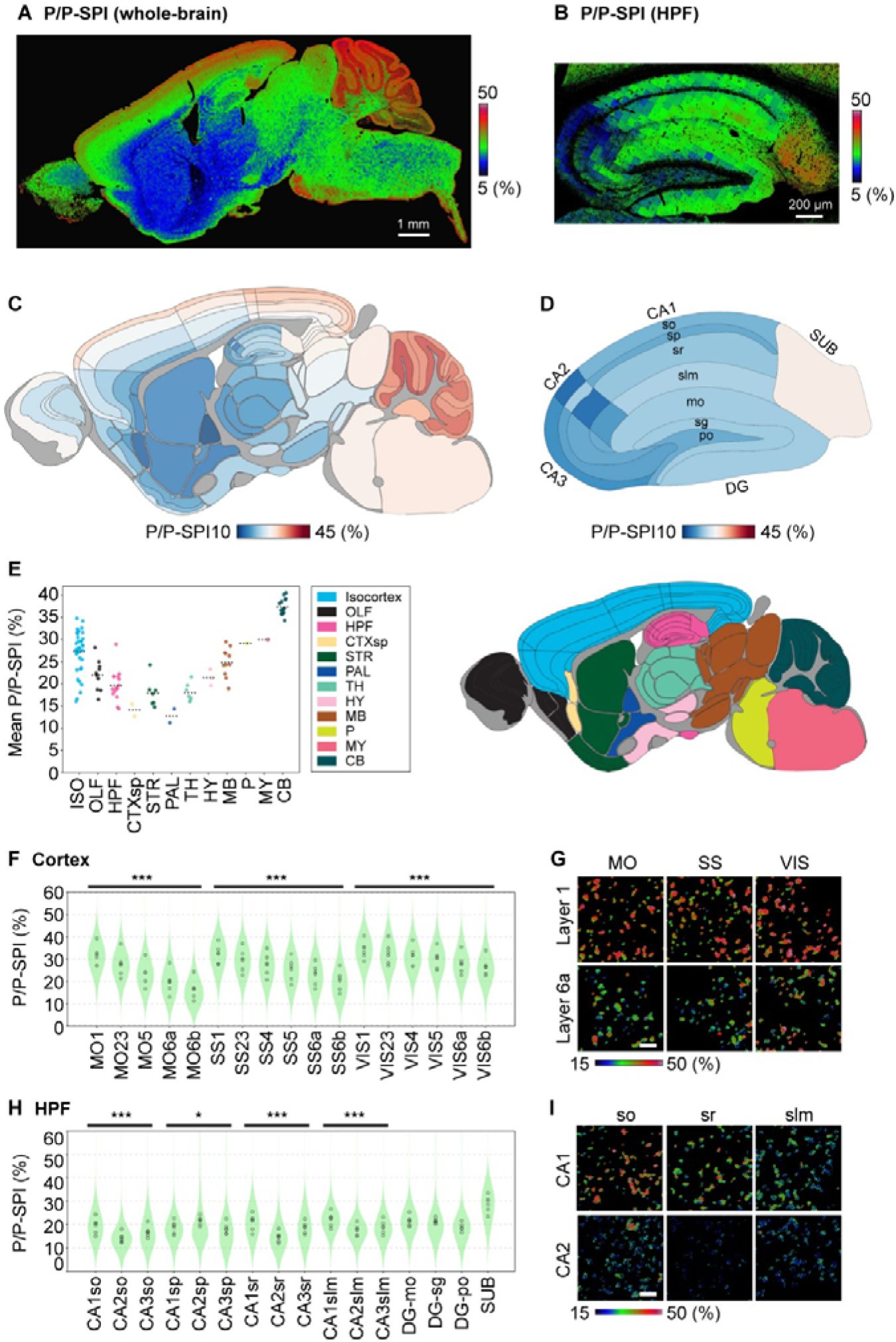
Association between PSD95 complexes differs across brain regions. (**A**,**B**) Representative example of P/P-SPI of individual tiles across whole brain (A) or HPF (B). (**C**,**D**) Heatmap of P/P-SPI of individual regions in whole brain (C) or HPF (D). The values are the average of six adult (4M) mice. (**E**) Dot plot of P/P-SPI in individual brain regions, grouped by overarching brain areas shown in the schematic (right). Black dotted lines indicate the mean for each area. The values are the average of six adult mice. For brain region abbreviations, see Supplementary Table 1. (**F**,**G**) Layer-dependent difference in P/P-SPI in cortical regions. Mean P/P-SPI of individual animals shown as dot plots and overall data distribution shown as violin plots behind (F). Representative examples of synapses in specific regions (G). MO: somatomotor areas; SS: somatosensory areas; VIS: visual areas. (**H**,**I**) Region-dependent difference in P/P-SPI in HPF. so: stratum oriens; sr: stratum radiatum; slm: stratum lacunosum-moleculare. *P < 0.05, ***P < 0.001, one-way ANOVA. Scale bars (G,I): 2 µm.

The highest levels of P/P-SPI were in superficial layers of the isocortex and cerebellum (Figs. 3C,E). In the isocortex, P/P-SPI values formed gradients along both the superficial–deep and anteroposterior axes (Figs. 3C,F,G). By contrast, the hippocampus, part of the allocortex, exhibited lower P/P-SPI than the more recently evolved isocortex (Figs. 3C,E). Within the hippocampus itself, different subregions, each with distinct roles in cognitive and social functions, also showed distinctive patterns of P/P-SPI diversity (Figs. 3D,H,I). The lowest P/P-SPI values were observed in subcortical structures including the striatum and thalamus, with the brainstem showing subregions with high values (Figs. 3C,E). Thus, synaptic PSD95 supercomplex proximity varies systematically across the brain, contributing to regional and circuit specialization. To facilitate access to this and other nanoscale synaptome atlases described below, we created the Mouse Synaptome Nanoarchitecture Brain Atlas Resource (URL https://git-pages.ecdf.ed.ac.uk/grantlab/nanoarchitecture-synaptome-atlas/).

### Lifespan trajectories in nanoscale synaptome architecture

The synaptome architecture of the brain, mapped through the synaptic composition of PSD95 and SAP102, undergoes continuous remodeling across the lifespan (lifespan synaptome architecture, LSA), with distinct features in each of three epochs (5, 6, 8, 37). The first epoch (LSA-1) spans from birth to adulthood (3M) and is marked by a dramatic expansion in the diversity and differentiation of the synaptome architecture of brain regions (5, 37). The second epoch (LSA-2), from 3M to 12M, is characterized by relative stability in synaptome architecture. The third epoch (LSA-3), from 12M onward, is marked by a loss of specific molecular types and subtypes of synapses, accompanied by a dedifferentiation of the synaptome architecture. Synaptome mapping of the rate of PSD95 turnover shows a progressive slowing across all LSA epochs and a preservation of synapses with the slowest turnover in LSA-3 (8).

Based on these previous observations, we proposed two hypotheses regarding temporal changes in the NSA: (i) that the P/P-SPI would increase during LSA-1 as protein complexes become more tightly packed into maturing synapses; and (ii) that this organization would remain stable throughout LSA-2 and LSA-3. To test these hypotheses, we used NanoSYNMAP to analyze 112 brain subregions in sagittal sections from PSD95–HaloTag mice at postnatal week 2 (2W), 4 (4M), 12 (12M), and 18 (18M) months of age (n = 6 animals per group). These data are incorporated into the Mouse Synaptome Nanoarchitecture Brain Atlas.

The brain heatmaps reveal a broadly similar spatial organization of P/P-SPI across all age groups (Fig. 4A). However, contrary to our initial predictions, we observed a widespread reduction in P/P-SPI across LSA-1 (2W to 4M), especially in the hippocampus, striatum, and thalamus. This indicates that very young animals have an abundance of synapses with tightly packed PSD95 supercomplexes, which declines as the brain matures (Fig. 4A-D). By contrast, during LSA-2 and LSA-3 (4M to 18M), there was a small gradual increase in P/P-SPI in most brain subregions (Fig. 4E).

**Figure 4.**
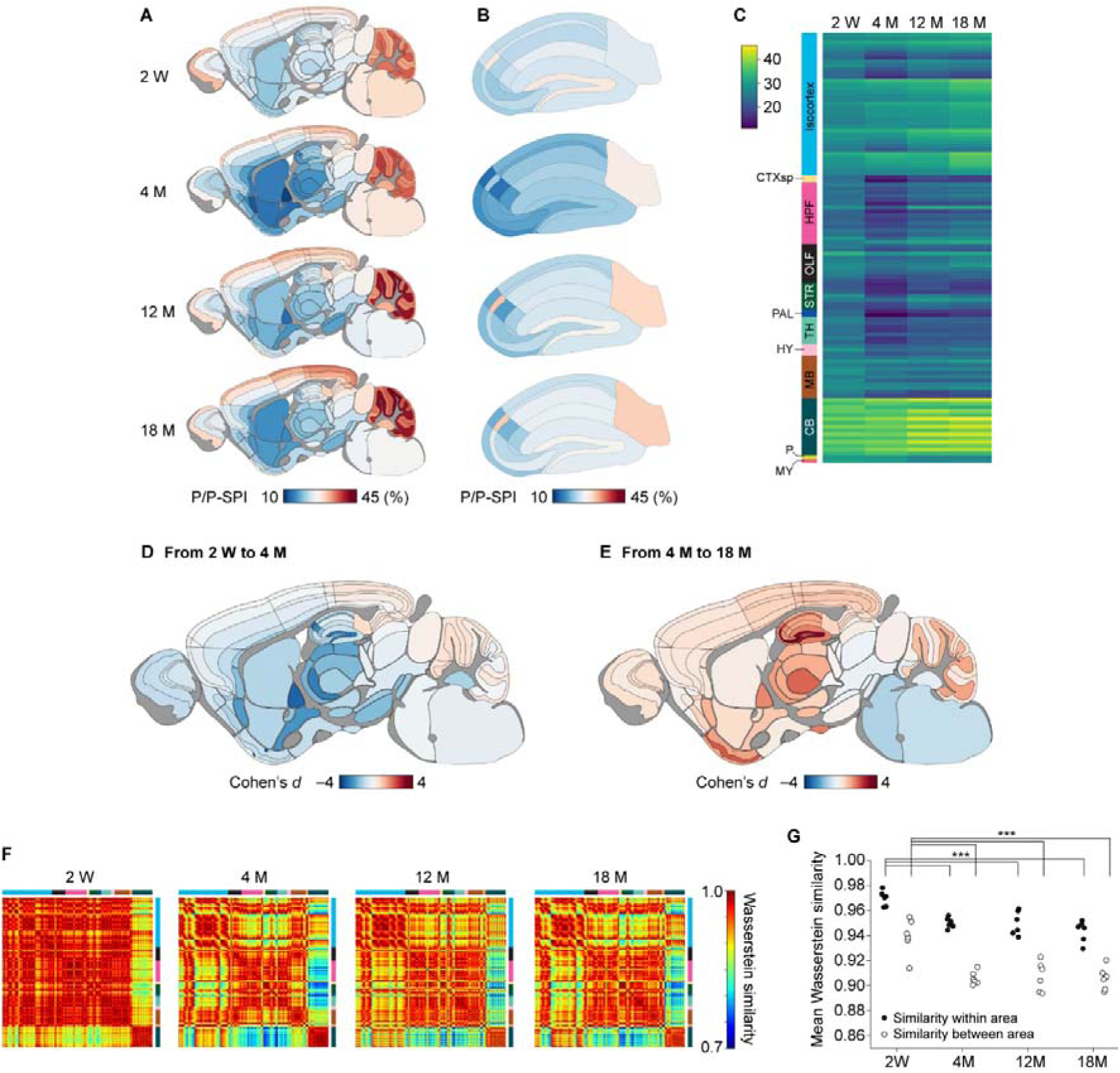
Lifespan trajectory of PSD95 complex association within synapses. (**A**,**B**) Heatmap of P/P-SPI of brain regions at different ages in whole brain (A) or HPF (B). (**C**) Integrated heatmap of average P/P-SPI of individual samples. (**D**,**E**) Cohen’s *d* of average P/P-SPI from 2W to 4M (D) or 4M to 18M (E). (**F**) Pairwise Wasserstein similarity matrices comparing P/P-SPI distributions between all subregion pairs at each age. Colored labels indicate overarching brain area annotations. **(G)** Per-animal summary of distribution similarity within and between brain areas across ages. For each animal, mean Wasserstein similarity was calculated across subregion pairs within the same brain area (filled circles) or between different brain areas (open circles) and plotted. ***P < 0.001, one-way ANOVA followed by Tukey’s post-hoc test.

To explore the possibility that the lifespan trajectories in the NSA affect the similarity between brain subregions, and thus the global synaptome architecture, we computed similarity matrices of the P/P-SPI distribution between all subregions at each age (Fig. 4F). In the youngest animals (2W), most brain subregions showed similarity in P/P-SPI, with the cerebellum again standing out as an outlier. By 4M, the subregions had differentiated into a distinct architecture, indicating that during the postnatal developmental period LSA-1 nanoscale organization contributes to regional differentiation. From 4M to 18M the similarity matrices are relatively stable. The differentiation of P/P-SPI during postnatal development was confirmed by quantification of the between-area and within-area similarity for each animal (Fig. 4G). Together, these results demonstrate that the NSA is regulated across the lifespan and contributes to the identity of individual brain subregions at different ages.

### Roles for protein abundance, clustering, and supercomplex competition in NSA diversity

To understand the mechanisms contributing to the proximity of supercomplexes within synapses, we modeled the arrangement of PSD95 beneath the postsynaptic membrane. We simulated a 500 nm diameter circular area populated with varying numbers of PSD95 molecules that were randomly distributed or clustered forming nanodomains (Fig. 5A). When randomly distributed, PSD95 copy number was linearly related to FRET efficiency, a FRET-based proximity metric corresponding to P/P-SPI (Fig. 5B). The introduction of nanodomains led to major increases in FRET efficiency (Fig. 5B). These simulations suggest that both the abundance of PSD95 and the presence of nanodomains are mechanisms that can contribute to the synaptic diversity in the P/P-SPI.

**Figure 5.**
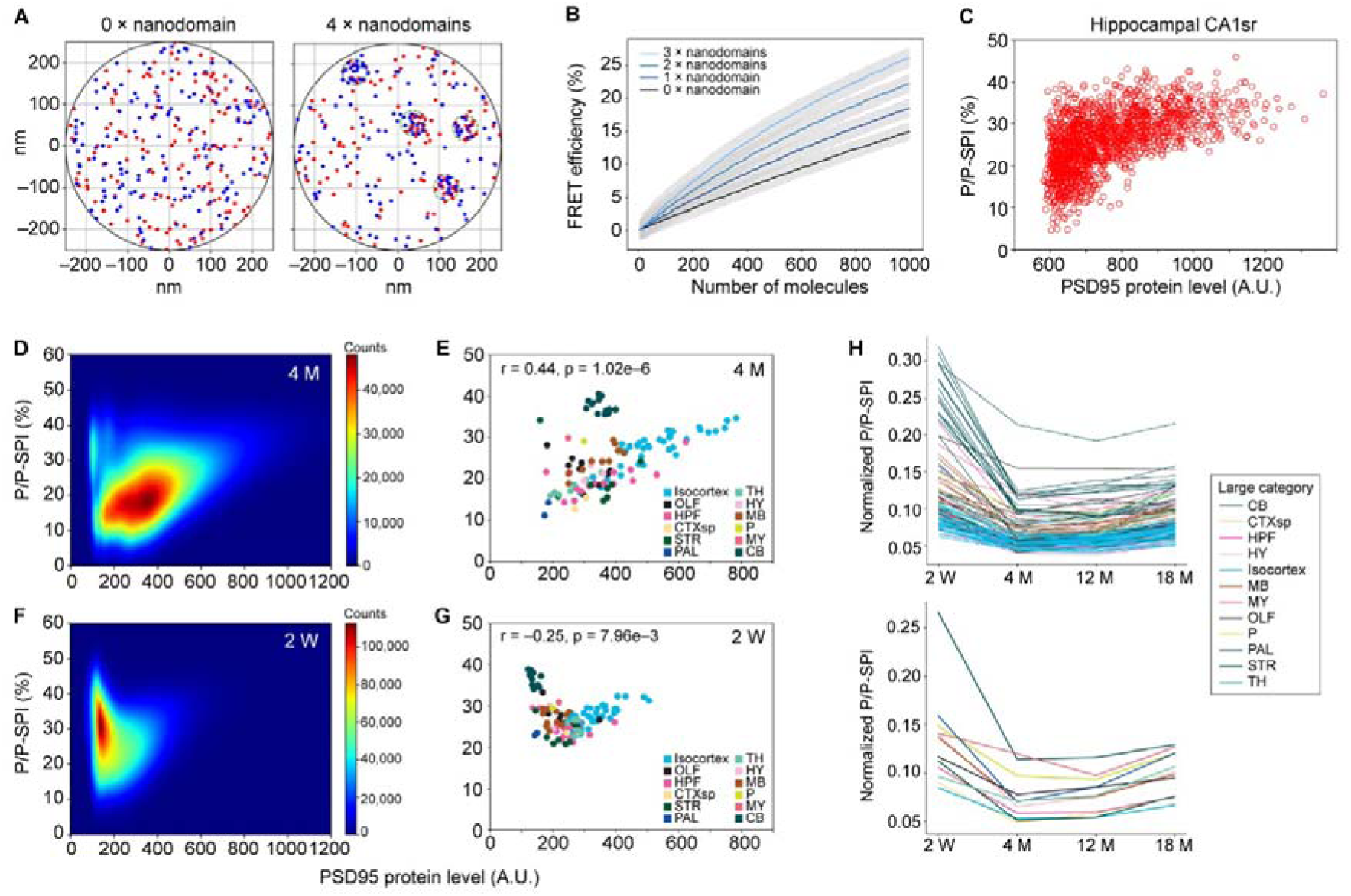
Relationship between PSD95 protein level and proximity of PSD95 supercomplexes. (**A**) Example of a random distribution of PSD95 molecules labeled with donor or acceptor on a synapse. In the simulation, 300 molecules (150 with donor and 150 with acceptor) were randomly distributed on the postsynaptic density, modeled as a circle of 500 nm diameter (left). Distribution in the presence of four nanodomains of 60 nm diameter (right). (**B**) Relationship between PSD95 density and FRET efficiency (corresponding to P/P-SPI). The mean P/P-SPI was obtained from 1,000 simulations assuming the indicated PSD95 copy number. Simulation was performed assuming 0, 1, 2, or 3 nanodomains. (**C**) Scatter plot of PSD95 protein level (acceptor fluorescence intensity) versus P/P-SPI for individual synaptic puncta in hippocampal CA1 region of adult (4M) PSD95_–_HaloTag mouse section. (**D-G**) Correlation between PSD95 protein level and P/P-SPI. (D,F) The data of all synaptic puncta of 4M or 2W mice are plotted as a 2D histogram heatmap. (E,G) Regional correlation between PSD95 protein level and P/P-SPI in 4M or 2W mice. Mean values of individual brain regions were plotted. (**H**) Plot of lifespan trajectory of normalized P/P-SPI (P/P-SPI divided by acceptor intensity) for individual brain regions.

To assess the relevance of these mechanisms *in vivo*, we plotted the P/P-SPI and PSD95 abundance (estimated from acceptor fluorescence) in CA1sr synapses (Fig. 5C). The spread in the observed values indicates that synapses vary in their nanodomain organization. Extending this analysis to the whole-brain synaptome datasets from 4M mice confirmed that these results apply across the brain (Fig. 5D). Representing the whole-brain data at the level of individual brain areas and subregions showed a positive correlation with protein abundance (r = 0.44, p = 1.02 × 10^-6^), with cerebellar subregions notably deviating from this trend (Fig. 5E). Excluding the cerebellar subregions from this analysis increased the correlation (r = 0.76, p = 1.10 × 10^-19^). By contrast, the positive correlation between PSD95 abundance and P/P-SPI was not observed at 2W at whole-puncta analysis (Fig. 5F) and in the brain region plot (r = -0.25, p = 7.96 × 10^-3^; Fig. 5G), but appeared at later ages (r = 0.393, p = 1.37 × 10^-5^; Figs. 5E and S4). This suggests that early postnatal synapses preferentially adopt nanoscale architectures that promote close PSD95 association, independent of total PSD95 abundance. Given the near-linear positive relationship between P/P-SPI and PSD95 abundance, we next sought to eliminate the contribution of total protein amount by normalizing P/P-SPI to PSD95 levels. Specifically, we divided P/P-SPI by the acceptor fluorescence intensity, yielding a PSD95-normalized P/P-SPI. Across most brain areas and subregions, this normalized metric exhibited a relatively stable pattern from adulthood (4M) onward, whereas a pronounced decrease was observed during postnatal development from 2W to 4M (Fig. 5H). This supports the idea that nanoscale postsynaptic architecture is actively reorganized during early life. Furthermore, the normalized P/P-SPI differed substantially across brain areas, with cerebellar subregions showing particularly distinctive values compared with the rest of the brain (Fig. 5H). Together with their deviation in the abundance–P/P-SPI relationship, this reinforces the conclusion that synaptic nanoscale organization is region-specific, and suggests that cerebellar synapses are characterized by especially close association of PSD95-containing complexes.

In our previous study, systematic imaging (at ∼240 nm resolution) of individual synapses expressing PSD95 and SAP102 across the whole mouse brain revealed that synapses differ in the amount, size, and shape of these proteins (4). These features permit classification into three excitatory synapse types based on protein expression (Type 1, PSD95 only; Type 2, SAP102 only; Type 3, PSD95 and SAP102) and 37 subtypes defined by quantitative differences in protein amount, size, and shape. Each type and subtype exhibits a distinct spatial distribution, forming the synaptome architecture of the brain. Having observed FRET between PSD95 and SAP102, we hypothesized that the presence of SAP102 supercomplexes around PSD95 supercomplexes could influence the proximity of PSD95 supercomplexes. To test this, we exploited the natural synaptic diversity in wild-type mice. Consistent with our hypothesis, correlations between P/P-SPI and the regional abundance of synapse subtypes revealed a clear distinction between Type 1 (which lack SAP102) and Type 3 (which contain SAP102) synapses (Fig. 6A). Synapses lacking SAP102 exhibited positive correlations, whereas those containing SAP102 showed negative correlations. Notably, Type 3 subtypes 20, 34, and 35 displayed weak positive correlations and these were the subtypes with the highest levels of PSD95. Individual plots of subtype abundance (subtypes 4 and 21) against P/P-SPI illustrated these respective positive and negative correlations (Fig. 6B). Further supporting our hypothesis, the proportion of Type 3 synapses in each brain subregion was strongly and negatively correlated with regional P/P-SPI (r = –0.77, p = 7.89 × 10⁻¹ ; Fig. 6C). These findings demonstrate that the presence of SAP102 interferes with the proximity of PSD95 supercomplexes.

**Figure 6.**
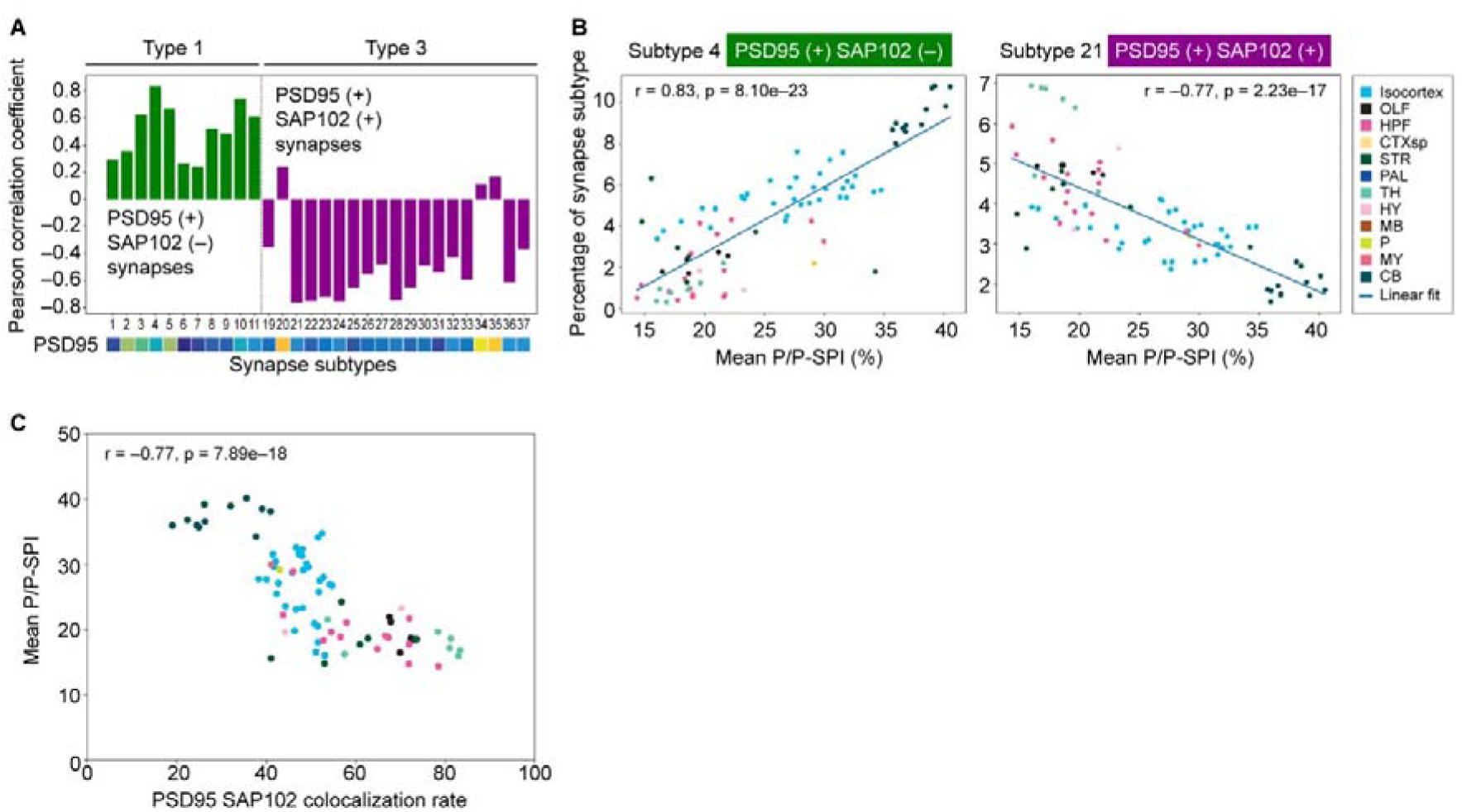
Correlation between PSD95 complex proximity and other synaptic parameters. (**A**) The correlation coefficient between mean P/P-SPI and ratio of individual synapse subtype population. (**B**) Representative examples of correlation between P/P-SPI and ratio of synapse subtypes that belong to Type 1 (subtype 4) or Type 3 (subtype 21). (**C**) Correlation between PSD95 SAP102 colocalization and P/P-SPI.

### PSD95 supercomplex abundance and proximity are regulated by PSD93

The above results suggest a general principle: combinations of supercomplexes govern the nanoscale spatial organization that underpins synaptome architecture. Ipso facto, the selective disruption of a supercomplex could rearrange postsynaptic density organization at the nanoscale, potentially leading to changes in the NSA. To test these predictions we employed *Dlg2^-/-^* mice, which lack PSD93 (11). *Dlg2^-/-^* mice show similar impairments in touchscreen cognitive tests as observed in schizophrenia patients carrying *DLG2* mutations (38). We crossed mice expressing PSD95-HaloTag with *Dlg2* knockout mice to generate *Dlg2^-/-^;Dlg4^HaloTag/HaloTag^*and controls (*Dlg4^HaloTag/HaloTag^*). Synaptome maps in 4M mice revealed a robust brainwide increase in P/P-SPI (Figs. 7A-D, S5A,B). The greatest increases were found in the isocortex (Fig. 7E), HPF (Fig. 7F) and other forebrain structures. The smallest P/P-SPI increases were in hindbrain structures, with the cerebellum showing the least increase. These data are incorporated into the Mouse Synaptome Nanoarchitecture Brain Atlas. Given our earlier finding that P/P-SPI is governed by both PSD95 abundance and nanodomain formation, we next asked whether the increase in P/P-SPI in *Dlg2^-/-^ ;Dlg4^HaloTag/HaloTag^* mice was driven by higher PSD95 levels. Correlation analysis of fold changes in P/P-SPI and PSD95 abundance revealed a positive relationship, as anticipated (r = 0.60, p = 2.40 × 10^-12^; Figs. 7G, S5C,D). However, in some regions (e.g. striatum), P/P-SPI increases exceeded those expected by protein abundance, whereas in others (e.g. hindbrain) P/P-SPI changed little despite elevated PSD95 levels. These findings indicate that loss of PSD93 supercomplexes leads not only to an increase in PSD95 supercomplex abundance, but also to altered proximity within synapses.

**Figure 7.**
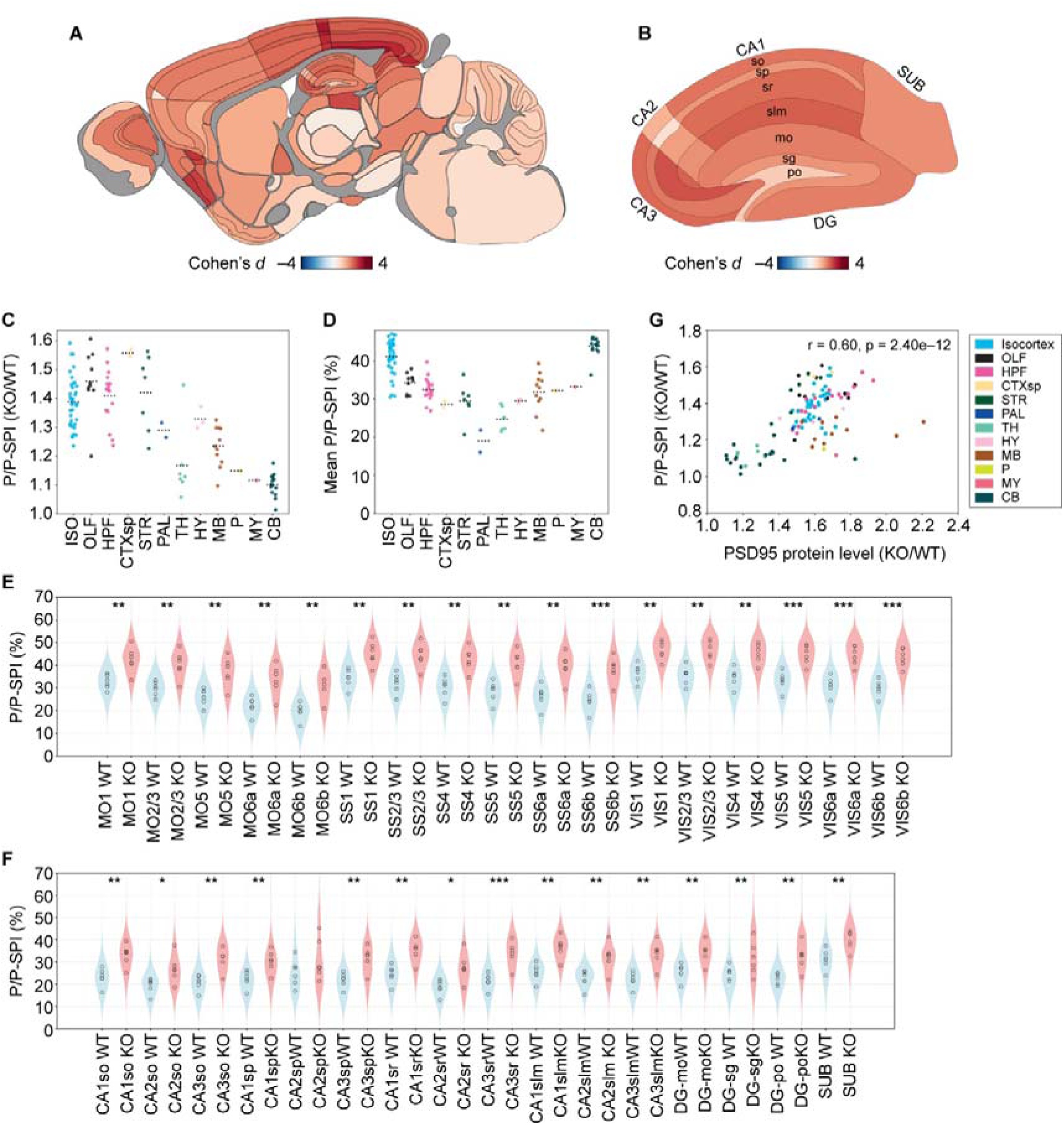
Lack of PSD93 results in reorganization of PSD95 complex association in synapses. (**A**,**B**) Heatmap of Cohen’s *d* of average P/P-SPI in whole brain (A) or HPF (B) of PSD93 knockout (*Dlg2*^-/-^, KO) (n = 6) compared with control (*Dlg2*^+/+^, WT) mice (n = 6). (**C,D**) Dot plots showing the fold change (KO/WT) of the average P/P-SPI in individual brain subregions (C) and the raw P/P-SPI values in individual brain subregions of PSD93 KO mice (D). Black dotted lines indicate the mean for each brain area. (**E**,**F**) Representative examples of the difference in P/P-SPI between WT and PSD93 KO mice in specific brain subregions. (**G**) Scatter plot showing fold change in mean P/P-SPI and PSD95 protein level between KO and WT across individual brain subregions. *P < 0.05, ** P < 0.01, *** P < 0.001; two-tailed unpaired Student’s t-test assuming equal variance.

### Relationship between PSD95 turnover and supercomplex proximity

A striking feature of adult mice lacking PSD93 is the reduced rate of PSD95 turnover and the corresponding increase in PSD95 protein lifetime (8). Together with the above observations, this suggests that synapses with more closely packed PSD95 supercomplexes may adopt a more stable configuration, reflected in longer protein lifetimes. To explore this possibility, we examined the relationship between PSD95 protein lifetime and PSD95 proximity by correlating their respective synaptome maps. In wild-type mice, we found that synapse subtypes with the longest PSD95 protein lifetimes (LPL synapses) showed strong positive correlations with P/P-SPI, whereas most subtypes with shorter lifetimes showed negative correlations (Fig. S6A). Across brain subregions, P/P-SPI and PSD95 half-life showed weak positive correlation (r = 0.25, p = 9.88 × 10^-3^; Fig. S6B), and this relationship was strengthened when analysis was restricted to isocortex and HPF (r = 0.72, p = 1.17 × 10^-9^). These results indicate that closer proximity of PSD95 supercomplexes is associated with slower protein turnover.

## DISCUSSION

A major challenge in neuroscience is to understand how proteins are organized at nanometer scales in individual synapses and how this organization is spatially and temporally regulated across the highly diverse and complex circuitry of the brain during the lifespan. Although super-resolution and ultrastructural methods have revealed nanoscale features of individual synapses, these approaches are not readily scalable to systematic, brainwide analyses at single-synapse resolution. Here, we developed NanoSYNMAP, a suite of genetic, optical, and computational tools that enables systematic evaluation of the nanoscale organization of synaptic proteins across the mouse brain. By integrating FRET as a molecular proximity reporter with synaptome mapping technology, we generated the Mouse Synaptome Nanoarchitecture Brain Atlas Resource, which contains brainwide, single-synapse resolution maps of postsynaptic MAGUK supercomplex proximity across the lifespan in mice. The spatial resolution of our approach is approximately 20-fold greater than previous mouse brain synaptome mapping approaches.

Our results reveal a previously unrecognized level of synaptic organization, which we term the nanoscale synaptome architecture (NSA). In this architecture, the proximity of MAGUK supercomplexes varies systematically across synapses, brain regions, and developmental stages. The NSA is distinct from synaptome architectures defined by protein composition, size, and shape at diffraction-limited resolution (4, 5, 37), and from ultrastructural descriptions of nanodomains within individual synapses (21, 22). Instead, the NSA captures relative molecular proximity within the postsynaptic proteome as a scalable and quantifiable dimension that can be mapped across millions of synapses.

Our results show that PSD95-containing and SAP102-containing supercomplexes can be positioned within several nanometers of one another, revealing that subsets of excitatory synapses contain tightly packed scaffold assemblies. The extent of this packing varies across synapse populations, generating nanoscale diversity that parallels the molecular and morphological diversity previously mapped at ∼240 nm resolution. The correlation between supercomplex proximity and PSD95 abundance indicates that local molecular density is a major determinant of nanoscale packing. However, systematic deviations from this relationship — most notably in the cerebellum — demonstrate that proximity cannot be explained by abundance alone. Cerebellar synapses exhibit unusually high PSD95 proximity despite relatively low PSD95 levels, consistent with ultrastructural evidence for dense, mesh-like postsynaptic densities and suggesting region-specific packing mechanisms (39–41). Furthermore, we found that the presence of nanoclusters is associated with higher proximity, consistent with these structures containing high densities of PSD95 supercomplexes. In this context, recent cryo-electron tomography studies describing clustered postsynaptic “nanoblocks” (42) provide a compelling structural correlate for the supercomplex proximities detected by NanoSYNMAP, supporting the idea that these nanoblocks correspond to MAGUK-based assemblies.

A key insight emerging from our findings is that nanoscale organization is governed by interactions between scaffold supercomplexes. The presence of SAP102- and PSD93-containing supercomplexes significantly influences PSD95 proximity, indicating that MAGUK assemblies interact competitively to shape nanoscale architecture. Genetic removal of PSD93 results in a robust, brainwide increase in PSD95 proximity and abundance, demonstrating that loss of one supercomplex species can drive nanoscale reorganization of others. These findings reveal an underlying molecular logic in which specific combinations of supercomplexes define local nanoscale architectures and, collectively, shape the global synaptome. The association of *DLG2* copy number variation with schizophrenia and autism (38, 43, 44) underscores the potential importance of nanoscale synaptome reorganization as a contributory factor in altered mental function.

Our findings in *Dlg2* mutant mice also provide mechanistic insight into compensatory processes that complicate interpretation of physiological and behavioral phenotypes in knockout models (45). Rather than just a loss of PSD93 function, we found that removal of this scaffold protein induced adaptive reorganization of nanoscale architecture, revealing the NSA as a flexible and adaptable system. This conclusion is supported by the observed changes in PSD95 abundance in *Dlg2* and *Dlg3* mutant mice measured using diffraction-limited synaptome mapping (4). The adaptability of the NSA is unlikely to be limited to mutations since exogenous and environmental influences, including sleep deprivation (7), head injury (10), and sensory deprivation (46), modify the synaptome architecture of PSD95 and SAP102. The capacity for nanoscale adaptation suggests potential therapeutic strategies that modulate synaptic protein proximity (47).

The lifespan atlas generated here demonstrate that the NSA is dynamically regulated with age. High PSD95 proximity in early postnatal brains indicates tightly packed supercomplex assemblies that separate during maturation and subsequently increase modestly during aging. Consistent with the competitive framework described above, a plausible contributor to the developmental decrease in PSD95 proximity is the progressive increase in PSD93 expression during maturation (11). The lifespan trajectory of the NSA parallels known phases of the LSA observed using diffraction-limited imaging (5). Both datasets reveal a dramatic global differentiation of brain subregions during the first 3 months of postnatal life followed by the transition to stable maintenance.

Integration of proximity synaptome maps with synapse protein lifetime maps indicates an association between nanoscale packing and proteostasis. Synapses and brain regions exhibiting higher proximity in adult mice show slower PSD95 turnover and reduced exchange of subunits within PSD95 dimers (20). These observations suggest that nanoscale organization is linked to molecular stability, with synapses enriched in superficial cortical layers exhibiting particularly stable supercomplex assemblies and slow protein turnover.

Nanoscale arrangements such as nanodomains and nanocolumns have been implicated in synaptic efficacy and plasticity (25–27), suggesting that proximity indices may relate to functional states. Postsynaptic scaffold proteins also play an important role in stabilizing AMPA receptors (48–50). Synapses with higher proximity may reflect more stable configurations optimized for reliable transmission, whereas those with lower proximity may be more permissive to remodeling and plasticity. Directly testing the relationship between nanoscale organization and synaptic function will require future studies combining measurements of protein composition and nanoarchitecture with single-synapse physiological recordings, which are not currently feasible.

There are several limitations of our approach. First, the proximities evaluated by this method are between fluorophores attached to genetically encoded C-terminal tags on endogenous MAGUKs. Second, FRET between multiple fluorophores provides an operational measure of molecular proximity rather than absolute distance, and proximity indices cannot be converted directly into nanometer-scale separations. Interpretation therefore relies on relative comparisons across regions, developmental stages, and genotypes. Future integration with super-resolution and ultrastructural techniques will further anchor proximity indices to physical organization.

The Mouse Synaptome Nanoarchitecture Brain Atlas Resource is designed to be extensible. While this study focuses on postsynaptic MAGUK scaffolds in excitatory synapses, the underlying framework can be readily applied to other postsynaptic and presynaptic proteins, including those in inhibitory and modulatory synapses. Our atlas can be readily integrated with transcriptomic, proteomic, connectomic, brain imaging and functional datasets, supporting multiscale models of brain organization.

## STAR★METHODS

### KEY RESOURCES TABLE

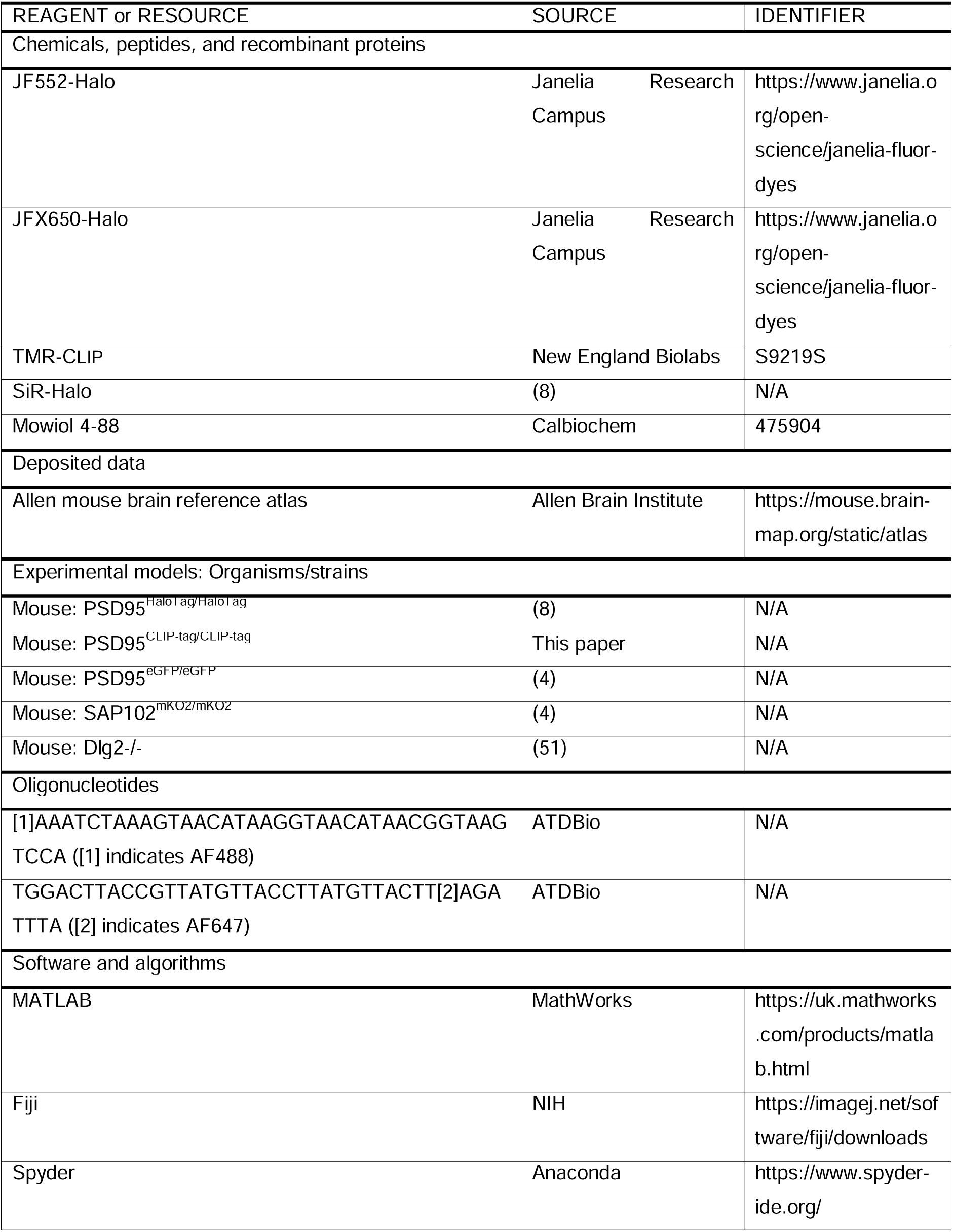

## EXPERIMENTAL MODEL DETAILS

### Mice

Animal experiments were approved by the Animal Welfare and Ethical Review Body (AWERB) of the University of Edinburgh. All animal procedures were carried out in accordance with the UK Animals (Scientific Procedures) Act 1986 and under a UK Home Office project license (PPL: PF3F251A9). C57BL/6J mice at the University of Edinburgh were used for the experiments. Mice were randomly selected with respect to sex, and approximately equal numbers of males and females were used overall.

## METHOD DETAILS

### Generation of a PSD95-CLIP-tag mouse line

First, a mouse line (PSD95 conditional CLIP-tag) was created using the previously reported method (52). This line harbors a conditional allele that enables the expression of the endogenous PSD95 protein with a CLIP-tag attached to its C-terminus in cells expressing Cre recombinase. In brief, an HDR template that removes an endogenous stop codon and inserts three stop codons flanked by two loxP sites, a short linker sequence (-SGGG-) and a CLIP-tag coding sequence in frame were integrated into the C-terminus of the mouse *Psd95* gene via CRISPR/Cas9-mediated genome editing in the zygote using the long single-stranded oligonucleotide method (52, 53). Then, this line was crossed with a CAG-Cre deleter line, which expresses Cre recombinase ubiquitously, including in germ cells, to create a mouse line (PSD95 constitutive CLIP-tag) that constitutively expresses the PSD95 protein with a CLIP-tag under the control of the endogenous regulatory elements. For genotyping, PCR was performed using genomic DNA. Primer F (5’-GTCACATGTCTTTGTGACCTTG -3’) and primer R1 (5’-GATACATGCAGAGAGGAGTGTC -3’) were used to amplify a 331 bp fragment of the wild-type allele. Primer F (5’-GTCACATGTCTTTGTGACCTTG - 3’) and primer R2 (5’-CTTCACCACTTTCAGCAGTTTC -3’) were used to amplify a 485 bp fragment of the knock-in allele.

### Tissue collection and section preparation

Mice were anesthetized by intraperitoneal injection of 0.1-0.2 ml (dependent on age) pentobarbital (Dolethal). A 26G needle was inserted into the left ventricle of the heart, and the right atrium was incised to allow outflow. The animal was transcardially perfused with 10 ml PBS followed by 10 ml fixative (4% paraformaldehyde (PFA)) and left at 4°C for 3 hours. Samples were transferred to 30% sucrose solution and incubated for 48-72 hours at 4°C. Brains were embedded in OCT compound (CellPath, KMA-0100-00A) in a plastic mold (Sigma-Aldrich). The molds in beakers containing isopentane (Sigma-Aldrich) were placed on liquid nitrogen for freezing. Frozen brains were stored at -80°C until use. Frozen brains were cut at 18 µm thickness using a cryostat (Leica CM3050 S) to obtain sagittal sections referring to bregma levels 12-14 from the Allen Mouse Brain Atlas. Cut brain sections were placed on Superfrost Plus glass slides (Epredia, J1800AMNZ) and dried at room temperature overnight in the dark and stored at 4°C.

### Postfixation labeling with HaloTag and CLIP-tag ligands

Brain sections were first incubated in PBS. 100 µl of the ligand solution in PBS was added to each brain section and incubated for 1 hour at room temperature in a wet dark chamber. For labeling of PSD95 (Halo/Halo) sections, Halo ligand conjugated with 666 nM JF552 and/or 1333 nM JFX650 was used. The combination of JF552 and JFX650 was selected based on preliminary evaluations, as it yielded the strongest FRET-derived fluorescence signal with low background among the available HaloTag ligands. For labeling of PSD95 (HaloTag/CLIP-tag) sections, CLIP ligand conjugated with 1 μM TMR and/or Halo ligand conjugated with 1 μM SiR was used. Brain sections were washed three times with PBS containing 0.2% Triton X-100 detergent and then twice with PBS for 10 minutes. Sections were then mounted using Mowiol 4-88 (Calbiochem, 475904) solution and covered with a coverslip (VWR, 631-0153).

### Spinning disk confocal microscopy

Imaging was performed using a Nikon spinning disk microscope (ECLIPSE Ti2) equipped with a 100x/1.4 NA objective lens (Nikon, MRD01902) with immersion oil (ZEISS, Immersol 518 F), as in our previous study (8). Images of 862 x 826 pixels in size and 16-bit depth were obtained with a Prime BSI Scientific CMOS Camera (Photometrics). To cover the whole area of the sagittal brain section, multi-tile single-plane image acquisition was performed. The following settings were used for individual channels. JF552 channel: excitation with 50 ms exposure of 561 nm laser at 1.91 mW; detection of fluorescence through BP 607/34 filter. JFX650 channel: excitation with 50 ms exposure of 640 nm laser at 5.53 mW; detection of fluorescence through BP 700/45 filter. FRET channel (JF552-JFX650): excitation with 200 ms exposure of 561 nm laser at 3.49 mW; detection of fluorescence through BP 700/45 filter. eGFP channel: excitation with 120 ms exposure of 488 nm laser at 1.06 mW; detection of fluorescence through BP 525/50 filter. mKO2 channel: excitation with 200 ms exposure of 561 nm laser at 2.33 mW; detection of fluorescence through BP 600/52 filter. FRET channel (eGFP-mKO2): excitation with 200 ms exposure of 488 nm laser at 4.00 mW; detection of fluorescence through BP 600/52 filter.

### Synapse detection and analysis

Synapse detection and FRET analysis were primarily conducted using custom Python scripts, with some downstream processing performed in MATLAB (54). For each image tile, synaptic structures were identified by independently applying Otsu’s method to both the donor and acceptor fluorescence channels, generating binary masks of fluorescence-positive regions. The overlapping areas between these two masks were extracted and defined as initial synaptic candidates. From these candidate regions, we excluded any area smaller than 5 pixels or larger than 5000 pixels, as well as regions containing saturated pixels, to reduce noise and potential artifacts. The remaining regions were designated as synaptic regions. For each synaptic region, the supercomplex proximity index (SPI) was calculated as FRET efficiency where the gamma factor is 1 (see “Calculation of supercomplex proximity index” for details). In the analysis of individual images, we generated TIFF images showing SPI values overlaid on synaptic masks, along with histograms and scatter plots to visualize the distribution of SPI across synapses. For brainwide synaptome analysis, numerical data — including SPI values for all detected synapses within each tile — were compiled into CSV files. To generate stitched whole-brain images at 1/16 scale, tile positions were extracted from the acquisition metadata, and tiles were assembled accordingly. The average synaptic values for each tile were overlaid onto the stitched image to create a brainwide heatmap. For region-of-interest (ROI) analyses, anatomical ROIs were manually defined using the 1/16-scale stitched images and the ROI Manager tool in Fiji. ROIs were delineated with reference to the sagittal atlas provided by the Allen Mouse Brain Atlas, and each ROI was saved as an individual .roi file. ROI labels were assigned according to the nomenclature of the bregma level 12 sagittal plate. When ROIs were delineated on the bregma level 13 or 14 plate, cerebellar ROI names were harmonized to the corresponding subregions on the bregma level 12 plate as follows: DEC was relabeled as SIM, FOTU as PRM and ANcr2, and PYR as COPY. Using these ROI definitions, tile-wise CSV files were converted into ROI-specific CSV files, aggregating synaptic data for each anatomical subregion. ROIs that could not be identified in at least three mice were excluded from subsequent analyses. Average SPI values were calculated per ROI and then averaged across animals to obtain group-level data. These results were visualized as dot plots, scatter plots, and heatmaps mapped onto a schematic of the sagittal brain layout.

### Calculation of supercomplex proximity index

For quantitative evaluation of FRET, we used a formula to calculate FRET efficiency from fluorescence intensity of donor and acceptor upon donor excitation (55). For simplicity, we calculated values in which gamma factor (56) is assumed as 1. These values were defined as SPI, which includes P/P-SPI and P/S-SPI. SPI was calculated as:

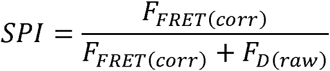

Where *F_FRET(corr)_* represents the corrected fluorescence intensity in the FRET channel (donor excitation, acceptor emission) and *F_D(raw)_* represents the fluorescence intensity in the donor channel. Corrections were applied to account for spectral bleed-through and direct excitation of the acceptor with the donor excitation laser.

The corrected signal in the FRET channel was calculated as:

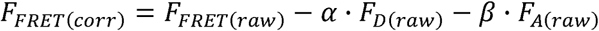

Here, *F_D(raw)_* and *F_A(raw)_* are the raw fluorescence intensities measured in the donor and acceptor channels. The coefficients and are correction factors accounting for donor bleed-through and acceptor direct excitation, respectively. The correction factors were calculated from control samples labeled with donor-only and acceptor-only fluorophores, as follows:

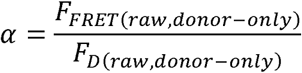

Where *F_FRET(raw,donor-only)_* and *F_D(raw,donor-only)_* are the raw fluorescence intensities in FRET and donor channels, respectively, measured in samples labeled with donor fluorophore.

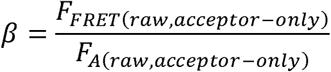

Where *F_FRET(raw,acceptor-only)_* and *F_A(raw,acceptor-only)_* are the raw fluorescence intensities in FRET and acceptor channels, respectively, measured in samples labeled with acceptor fluorophore.

For quantification of raw fluorescence intensity in individual channels, background subtraction was performed. This method allowed us to quantitatively estimate SPI on a synapse-by-synapse basis while correcting for channel crosstalk and optical artifacts.

### Analysis of data distributions and similarities across brain regions

To characterize the distribution of SPI values within each anatomical region, we aggregated ROI-level datasets across animals within each experimental group using custom Python scripts. For each group and ROI, SPI values from all contributing animals were pooled. The pooled values were then converted into a binned distribution with 100 equally spaced bins spanning 0.00–1.00. For each bin, the percentage of synapses falling into the bin was computed relative to the total number of synapses in that ROI and group. To quantify similarity between regional SPI distributions, we computed pairwise Wasserstein similarity matrices from the binned distributions. The 100-bin percentage vectors were normalized to sum to 1. For each group, the 1D Wasserstein distance between all pairs of ROI distributions was calculated from the mean absolute difference between their cumulative distribution functions over the 100 bins. Wasserstein similarity was then defined as 1 − Wasserstein distance, yielding values approaching 1 for highly similar distributions. Similarity matrices were visualized as heatmaps with ROI order defined by a curated ROI list, and ROIs were annotated by overarching brain area classification. In addition, we quantified similarity across brain regions at the level of individual animals. For each animal and ROI, SPI values were binned into 100 equally spaced bins. Wasserstein similarity was then computed for all ROI pairs as described above. For each animal, mean within-area similarity and mean between-area similarity were computed across all eligible ROI pairs and used as summary measures of regional similarity.

### Statistical analysis

For the analysis of distribution similarity, Wasserstein similarity values were used as per-animal summary metrics and compared across age groups using one-way ANOVA, followed (when appropriate) by Tukey’s honestly significant difference (HSD) test for post-hoc pairwise comparisons.

Cohen’s *d* values that measure the effect size of synaptome parameter changes between two groups of mice were calculated as described previously (6):

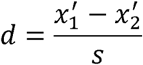

where 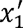 and 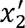 are the sample average and s is pooled standard deviation as follows:

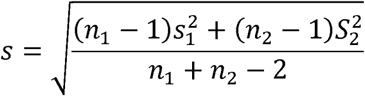

Where *s*_1_ and *s*_2_ are the sample standard deviation and *n*_1_ and *n*_1_ are the sample numbers of individual groups.

Statistical comparisons between two groups were performed using two-tailed t-tests implemented with the T.TEST function in Microsoft Excel.

### Preparation of brain extracts for single-molecule confocal microscopy

Preparation of brain extracts was performed using a previously described method with minor modifications (57). Half hemispheres from PSD95–HaloTag mice were lysed in 2 ml lysis buffer (1% deoxycholate, 50 mM Tris-HCl pH 9.0, 15 mM NaF, 20 µM ZnCl_2_, 2.5 mM sodium orthovanadate, and cOmplete EDTA-free Protease Inhibitor Cocktail (Roche)). Homogenization was performed using 20 strokes in a Potter Homogenizer. The homogenates were incubated on ice for 1 hour and then centrifuged for 30 minutes at 50,000 g using an Optima MAX-XP ultracentrifuge. The supernatants were taken and stored at -80°C until use. For HaloTag labeling, a 1:1 mixture of HaloTag ligands conjugated with AF488 and JFX650 was added to the lysates. The final reaction mixture contained 10 mg/ml total protein and 200 nM of each ligand and was incubated on ice for 1 hour. After labeling, the samples were diluted 40,000-fold with PBS. As a positive control, we used a dual-labeled DNA sample with two fluorophores positioned 2 nm apart (ATDBio). The oligonucleotide sequences are as follows: 5’-[1]AAATCTAAAGTAACATAAGGTAACATAACGGTAAGTCCA-3’ and 5’-TGGACTTACCGTTATGTTACCTTATGTTACTT[2]AGATTTA-3’, where [1] indicates AF488 and [2] indicates AF647.

### Preparation of microfluidic device

The microfluidic device design has been described previously (58). It consists of a single microchannel (width 100 μm, height 25 μm, length 1 cm). Devices were fabricated using standard soft-lithography techniques, in which polydimethylsiloxane (PDMS; Dow Corning) was molded on SU-8 photoresist-patterned silicon wafers, as described previously. The PDMS devices were bonded to glass coverslips (VWR, thickness No. 1) using oxygen plasma treatment (ZEPTO B, Diener) to form sealed microchannels.

### Single-molecule confocal measurements

All confocal microscopy measurements were performed using a custom-built single-molecule confocal microscope described previously (59, 60). Excitation was provided by a laser combiner (Oxxius L6Cc, Oxxius) equipped with 488 nm and 638 nm Gaussian lasers. The beams were coupled into a 0.13 NA optical fiber (Thorlabs), collimated and introduced through the back-port of an inverted microscope (Nikon TE2000-U) via a reflective collimator (Thorlabs). Excitation was focused to a confocal volume using a 100x oil-immersion objective lens (Nikon CFI Plan Apochromat VC, NA 1.4). The emitted fluorescence was collected through the same objective lens and passed through a primary dichroic mirror (DI03-R405/488/561/635, Semrock), and a 50 μm pinhole (Thorlabs). A secondary dichroic mirror (585DRLP, Omega Filters) separated the emission signals by wavelength. Fluorescence from the far-red and green channels was directed through appropriate long-pass and band-pass filters (green: BLP01-488R-25 and FF01-525/30-25; far-red: LP02-647RU-25 and FF01-432/515/595/730-25; Semrock) and detected using two independent avalanche photodiodes (PerkinElmer). Photon counts were recorded using a USB data acquisition card (USB-CTR04, Measurement Computing), with 100 μs time bins, the expected residence time of the PSD95 complexes in the confocal volume. Laser intensities at the back port were 0.604 mW (488 nm) and 0.395 mW (638 nm). Samples were introduced into the microfluidic device using a 200 µl gel-loading tip. Flow was maintained using a negative-pressure-driven syringe (Injekt®-F Luer Solo, B.Braun) controlled by a precision syringe pump (LEGATO 101, World Precision Instruments) at a flow rate of 96 µl/h. Each sample was measured for 15 minutes. Data analysis was performed using custom scripts written in Python. Coincident fluorescence events were defined as events with at least 10 photon counts per 100 µs bin in both channels. The coincidence fraction was calculated as the ratio of coincident events to the total number of events with above 10 photons per bin in the AF488 channel.

### Simulation of FRET efficiency

FRET simulations were performed using custom Python scripts. In the simulation, a synapse was modeled as a two-dimensional circular area with a diameter of 500 nm. An even number of point particles (representing PSD95 molecules) were randomly distributed within this circular region. Half of the particles were randomly designated as donor-labeled molecules, and the remaining half as acceptor-labeled molecules. For each donor molecule, the distance to the nearest acceptor was calculated, and the FRET efficiency was estimated based on the Förster equation:

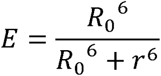

where *r* is the distance between the donor and the nearest acceptor, and *R*_0_ is the Förster radius, set to 4.2 nm in the simulation. The mean FRET efficiency across all donor molecules within a single synapse was calculated and used as the representative FRET efficiency of that synapse. To investigate the relationship between the number of PSD95 molecules and synaptic FRET efficiency, simulations were performed with PSD95 counts ranging from 2 to 1000 (in even-numbered increments). For each condition, 1000 independent simulations were conducted, and the mean ± standard deviation of FRET efficiency was plotted. To simulate the formation of nanodomains, 0, 1, 2, or 3 circular subregions with a diameter of 60 nm were introduced within the synaptic area. Each subregion contained 10% of the total number of PSD95 molecules, while the remaining molecules were randomly distributed throughout the synapse.

### Estimation of Förster radius

The Förster radius (R₀) for the JF552–JFX650 donor–acceptor pair was calculated using the following equation (31):

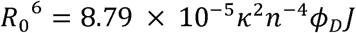

The orientation factor (κ²) was assumed to be 2/3, consistent with the dynamic isotropic averaging model. The refractive index (n) of Mowiol 4-88 and the fluorescence quantum yield (Φ_D_) of JF552 were set to 1.381 and 0.83, respectively, based on the supplier’s specifications. The spectral overlap integral (J) between the JF552 emission spectrum and the JFX650 absorption spectrum was estimated as 4.2 × 10¹, calculated from the supplier-provided spectral data. For quantitative extraction of spectral values, plots were digitized using WebPlotDigitizer (https://automeris.io/). The spectral overlap integral was then computed using the equation:

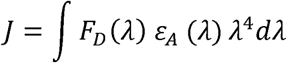

where *F_D_(λ*) is the donor emission spectrum normalized such that 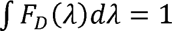, *ε_A_(λ*) is the acceptor molar extinction coefficient at wavelength λ, and λ is the wavelength (nm). Accordingly, J is expressed in units of M^-1^ cm^3^ nm^4^.

## Code availability

Scripts for the core processing steps of the FRET synaptome mapping pipeline used in this study are available at GitHub (https://github.com/Takeshi-Kaizuka/Kaizuka_NanoSYNMAP_2026) and Zenodo (https://doi.org/10.5281/zenodo.18486629) (54).

## ACKNOWLEDGEMENTS

We thank Hanan Woods for providing the analytical dataset, Emily Robson for the management of mouse lines, Ulku Gunar, Vilhelmiina Savolainen, Theresa Wong, and Beverley Notman for technical assistance, and Colin Davey for the critical reading of the manuscript. T.K. acknowledges funding from Uehara Memorial Foundation Research Fellowship and the European Union’s Horizon 2020 Research and Innovation Programme under Marie Sklodowska-Curie grant agreement No 101029343 (SYNarch). T.Z. was supported by a Centre for Doctoral Training in Tissue Repair, Innovation and Collaboration (CenTRIC) studentship funded by the Eureka Foundation. C.A. acknowledges support from the BBSRC EastBIO Doctoral Training Programme (BB/M010996/1). Work in the S.G.N.G. lab was funded by the Wellcome Trust (302077/Z/23/Z, 221295/Z/20/Z), the European Research Council (ERC) under the European Union’s Horizon 2020 Research and Innovation Programme (885069 SYNAPTOME) and Simons Initiative for the Developing Brain (SIDB) under the Simons Foundation for Autism Research Initiative (529085). The single-molecule confocal microscope used in this study was funded by Alzheimer’s Research UK (ARUK-EG2018B-004) and a kind donation from Dr Jim Love. For the purpose of open access, the author has applied a CC-BY public copyright licence to any Author Accepted Manuscript version arising from this submission.

## AUTHOR CONTRIBUTIONS

T.K., M.H.H., and S.G.N.G. organized the project. T.K., E.B., and C.A. planned and performed imaging of brain sections and synaptosomes. T.K., Z.Q., D.D., and M.H.H. developed the computational pipeline for image processing and data analyses. T.K., K.M., T.Z., and M.H.H. performed single-molecule confocal microscopy. G.V. and N.H.K. developed PSD95-CLIP mouse line. T.K. and S.G.N.G. wrote the manuscript. M.H.H. and S.G.N.G. supervised the project. All authors reviewed the manuscript and approved the final version.

## SUPPLEMENTAL INFORMATION

**Figure S1.**
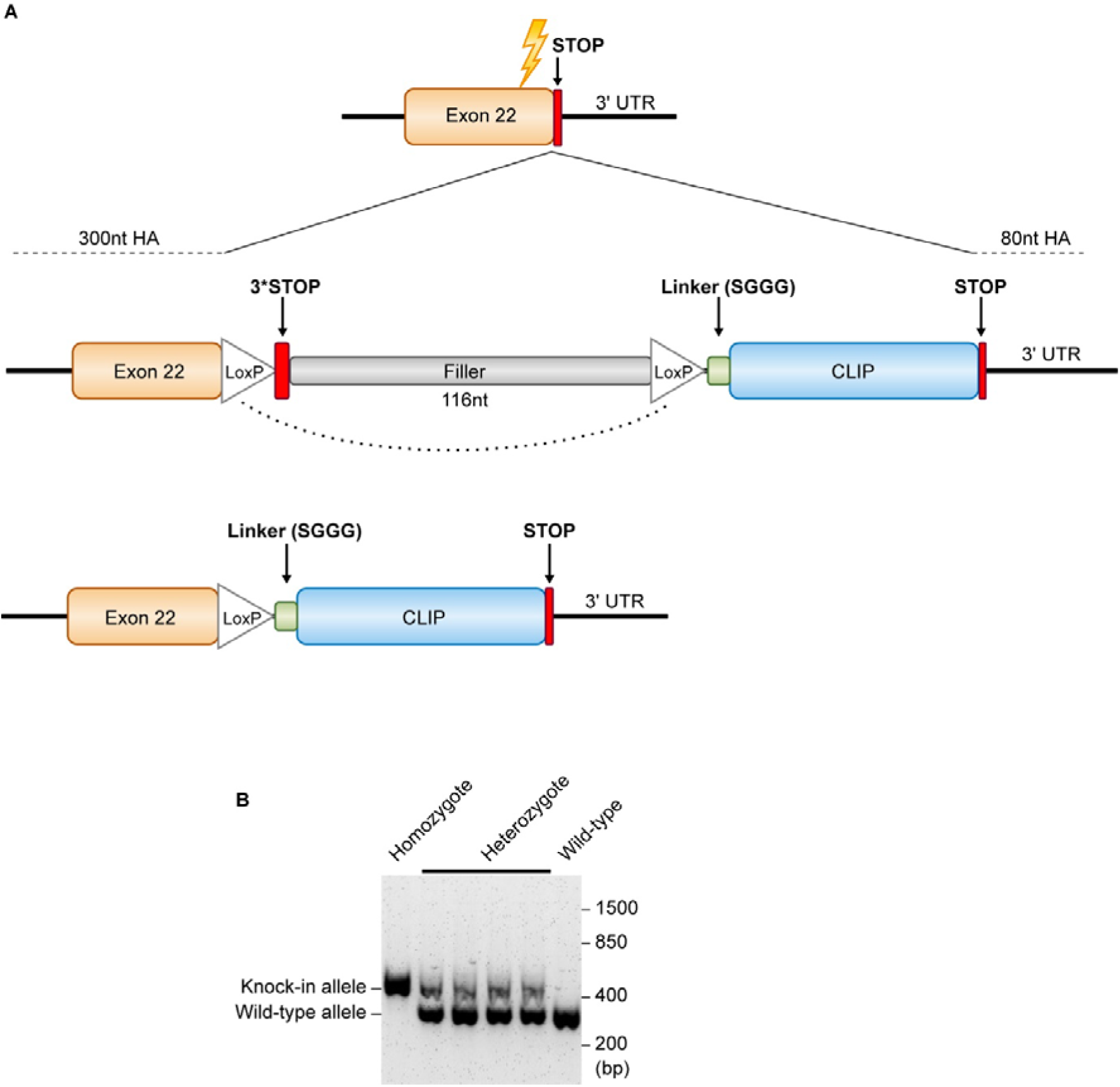
Generation of knock-in mice that express PSD95 tagged with CLIP-tag. (**A**) Targeting strategy for the *Psd95* (*Dlg4*) genomic locus. The *Dlg4* allele was targeted by inserting the CLIP-tag coding sequence at the end of exon 22, together with loxP sites and a filler sequence, to generate a conditional knock-in line. The sequence between the two loxP sites was then excised by crossing the conditional knock-in line with a Cre recombinase–expressing transgenic mouse line. (**B**) Representative genotyping PCR. PCR amplification distinguishes the wild-type and knock-in alleles: the wild-type allele yields a 331 bp fragment, whereas the knock-in allele yields a 485 bp fragment.

**Figure S2.**
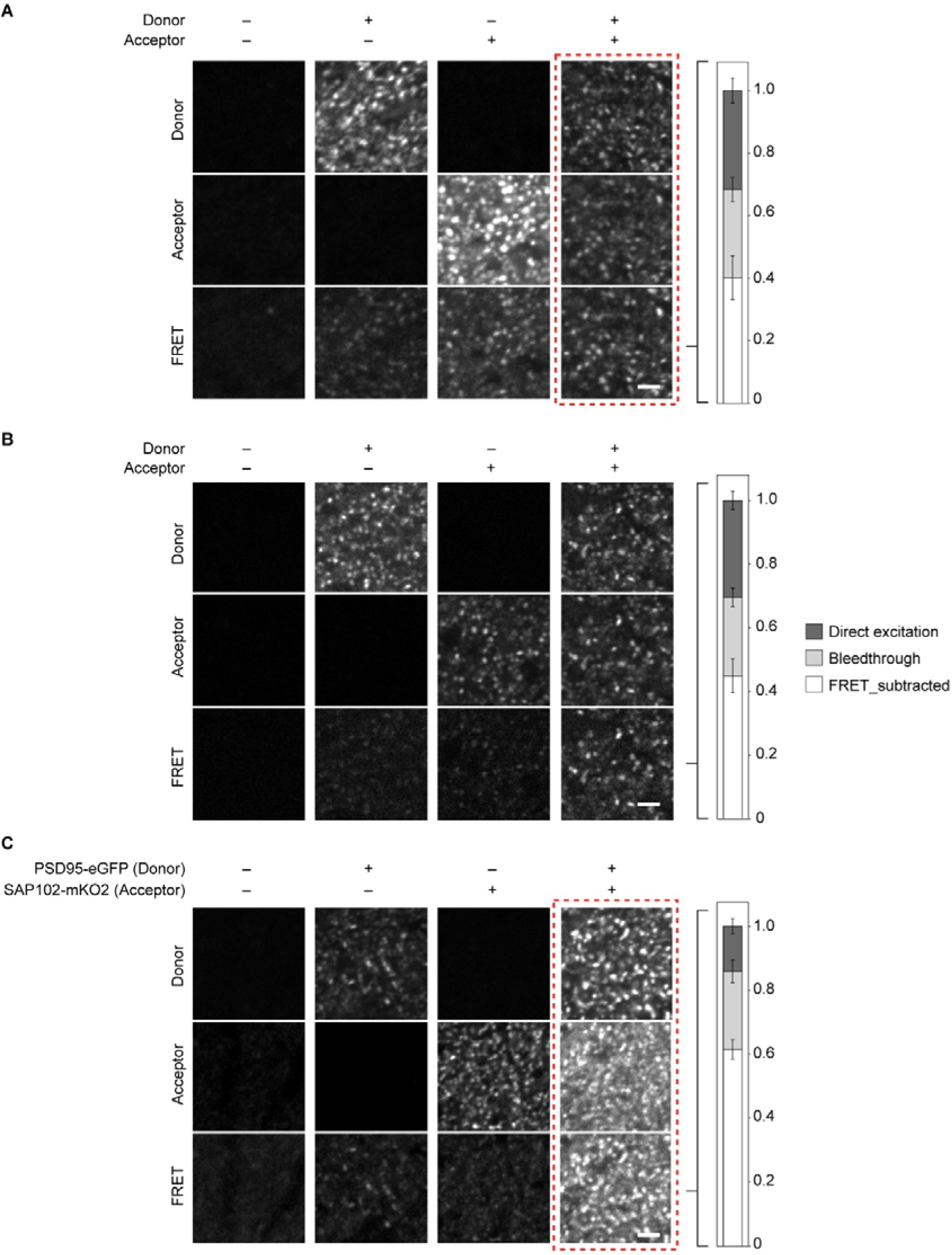
Detection of fluorescence and FRET in hippocampal CA1 region of the mouse brain. (**A**) Sections obtained from PSD95 (Halo/Halo) mouse were labeled with HaloTag ligands conjugated with donor, acceptor, or their mixture. The fluorescence of the donor, acceptor, and FRET between them was detected by confocal microscopy. The proportions of fluorescence in FRET channel attributable to FRET, bleedthrough (cross-talk) and direct excitation (of acceptor fluorophore) were estimated and plotted (right). Boxed panels are shown in Fig. 1D. (**B**) Sections obtained from PSD95 (HaloTag/CLIP-tag) mouse were labeled with CLIP-tag ligand (donor) and HaloTag ligand (acceptor) conjugated with the respective fluorophores. The fluorescence of the donor, acceptor, and FRET between them was detected by confocal microscopy. The proportions of fluorescence in FRET channel attributable to FRET, bleedthrough and direct excitation were estimated and plotted (right). (**C**) Sections obtained from PSD95 (eGFP/eGFP) SAP102 (mKO2/mKO2) mouse or mouse expressing either PSD95-eGFP or SAP102-mKO2 were prepared. The fluorescence of the donor, acceptor, and FRET between them was detected by confocal microscopy. The proportions of fluorescence in FRET channel attributable to FRET, bleedthrough and direct excitation were estimated and plotted (right). Boxed panels are shown in Fig. 1F. Scale bars: 2 µm.

**Figure S3.**
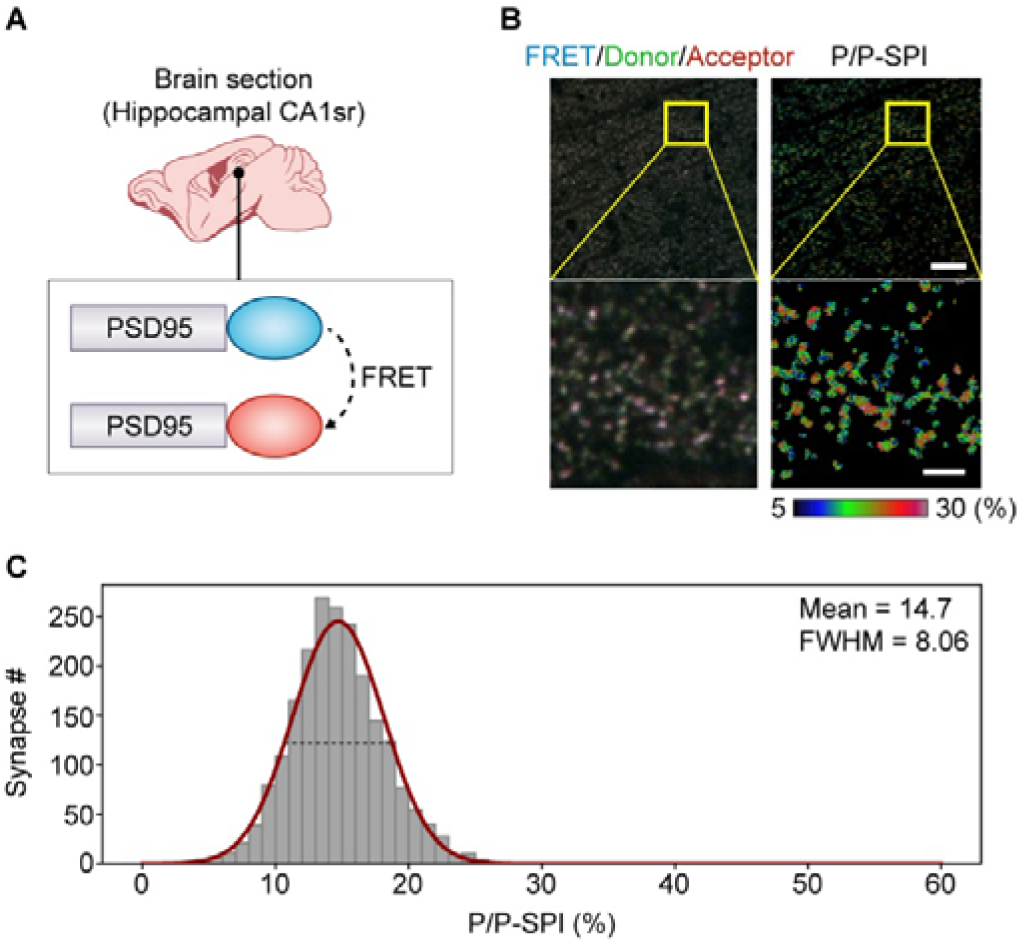
Diversity of synaptic nanoarchitecture revealed by FRET analysis in PSD95–HaloTag/CLIP-tag mouse. (**A**) Visualization of synaptic P/P-SPI in the hippocampal CA1sr region using PSD95–HaloTag/CLIP-tag mouse. Sections obtained from PSD95 HaloTag/CLIP-tag mouse were labeled with a mixture of HaloTag ligand conjugated with acceptor and CLIP-tag ligand conjugated with donor and observed by confocal microscopy. (**B**) (Left) Merged image of JF552, JFX650, and FRET channels. (Right) P/P-SPI of individual synaptic puncta. Scale bars: 10 µm (top) and 2 µm (bottom). (**C**) Histogram of P/P-SPI of individual synaptic puncta. The Gaussian fit to the distribution (red line), together with its mean value and full width at half maximum (FWHM; black dashed line), are shown.

**Figure S4.**
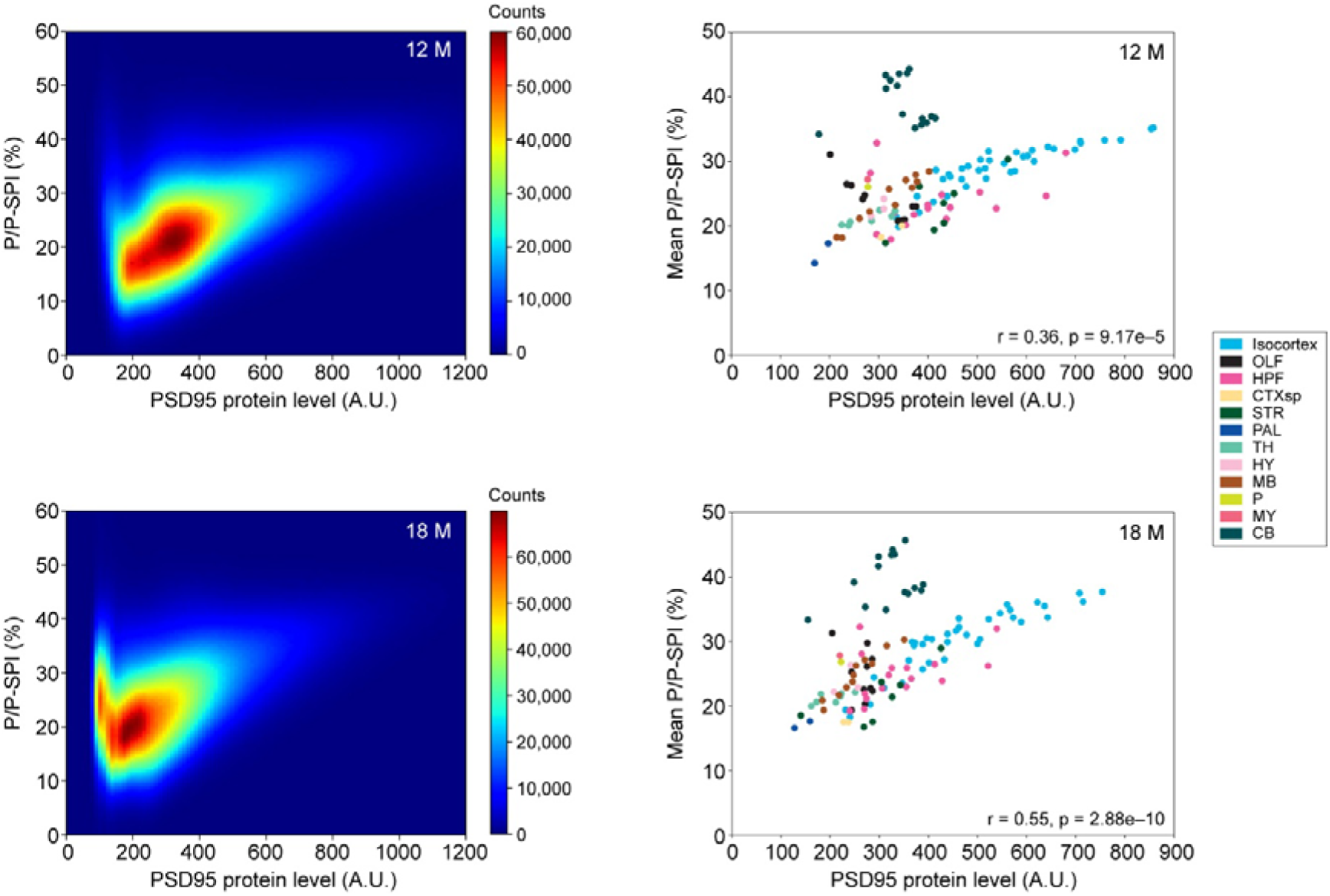
Relationship between PSD95 protein level and P/P-SPI in aged mice. Correlation between PSD95 protein level and P/P-SPI (related to Fig. 5). (Left) Data from all synaptic puncta in 12M (top) and 18M (bottom) mice are shown as a 2D histogram heatmap. (Right) Regional correlations between PSD95 protein level and P/P-SPI in 12M (top) and 18M (bottom) mice.

**Figure S5.**
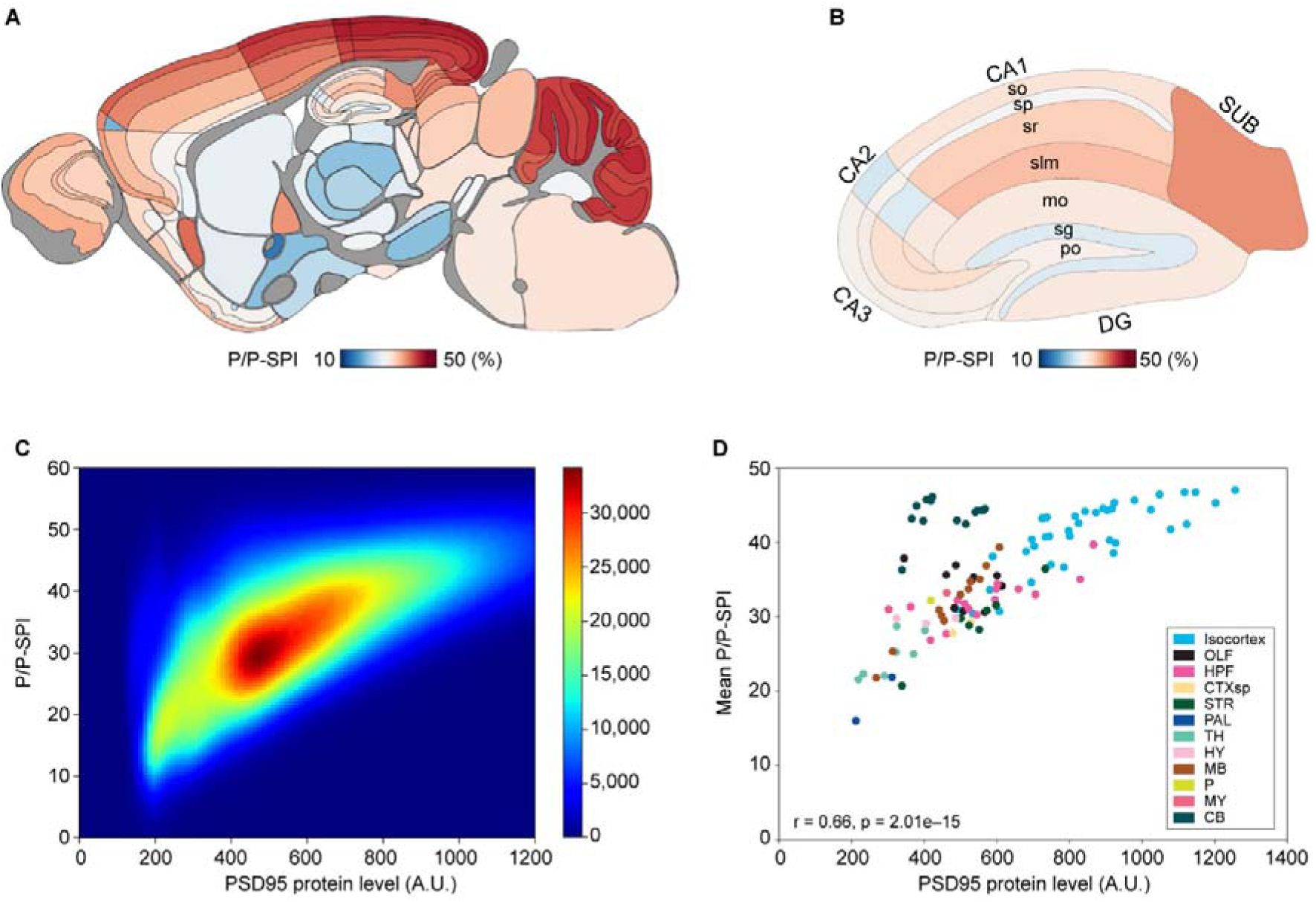
Overview of FRET synaptome analysis of PSD93 KO mice. (**A**,**B**) Heatmap of P/P-SPI of individual subregions in whole brain (A) or hippocampus (B). Values are the average of six PSD93 KO mice. (**C**) Correlation between PSD95 protein level and P/P-SPI. The data of all synaptic puncta of PSD93 KO mice are plotted as 2D histogram heatmap. (**D**) Regional correlation between PSD95 protein level and P/P-SPI in PSD93 KO mice.

**Figure S6.**
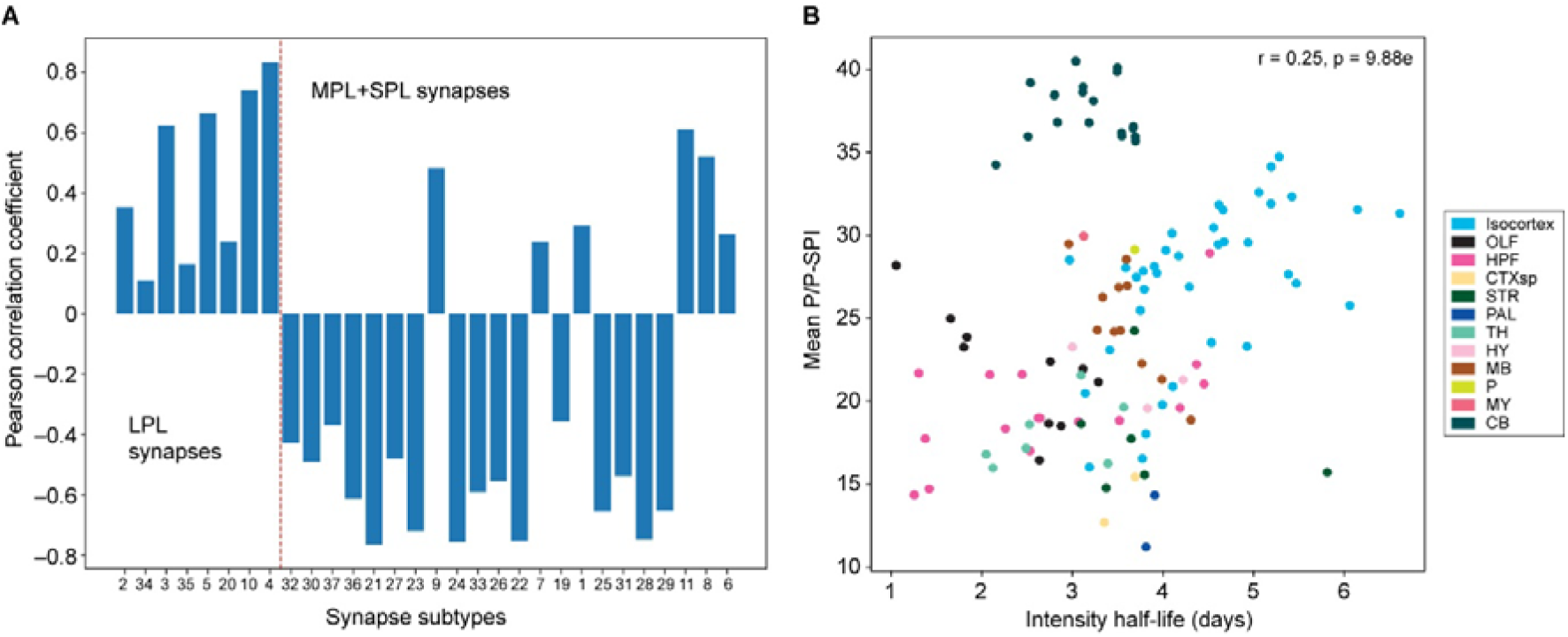
Correlation between P/P-SPI and PSD95 protein lifetime. (**A**) The correlation coefficient between mean P/P-SPI and ratio of individual synapse subtype population. The plotted data are identical to Fig. 6A but here the synapse subtypes are ordered based on their PSD95 protein lifetime as shown in our previous study (8). (**B**) Scatter plot of regional P/P-SPI measured in 4-month-old mice and PSD95 protein half-life determined in 3-month-old mice (8).

**Supplementary Table 1. List of brain regions analyzed in this study**

List of the 112 brain regions analyzed in the present study. For each region, the table provides the abbreviation and full anatomical name, together with the corresponding brain area (abbreviation and full name) to which the region is assigned. All nomenclature follows the Allen Brain Atlas ontology.

